# Suppression of tumor cell lactate-generating signaling pathways eradicates murine PTEN/p53-deficient aggressive-variant prostate cancer via macrophage phagocytosis

**DOI:** 10.1101/2023.05.23.540590

**Authors:** Kiranj Chaudagar, Hanna M. Hieromnimon, Anne Kelley, Brian Labadie, Jordan Shafran, Srikrishnan Rameshbabu, Catherine Drovetsky, Kaela Bynoe, Ani Solanki, Erica Markiewicz, Xiaobing Fan, Massimo Loda, Akash Patnaik

**Author notes:** **Corresponding Author:** Akash Patnaik, M.D., Ph.D., M.M.Sc. Knapp Center for Biomedical Discovery, 7152 900 East 57th Street Chicago, IL 60637.

## Abstract

**Purpose:** PTEN loss-of-function/PI3K pathway hyperactivation occurs in ∼50% of metastatic, castrate-resistant prostate cancer patients, resulting in poor therapeutic outcomes and resistance to immune checkpoint inhibitors across multiple malignancies. Our prior studies in prostate-specific PTEN/p53-deleted genetically engineered mice (Pb-Cre;PTEN^fl/fl^Trp53^fl/fl^ GEM) with aggressive-variant prostate cancer (AVPC) demonstrated feedback Wnt/β-catenin signaling activation in 40% mice resistant to androgen deprivation therapy (ADT)/PI3K inhibitor (PI3Ki)/PD-1 antibody (aPD-1) combination, resulting in restoration of lactate cross-talk between tumor-cells and tumor-associated macrophages (TAM), histone lactylation (H3K18lac) and phagocytic suppression within TAM. Here, we targeted immunometabolic mechanism(s) of resistance to ADT/PI3Ki/aPD-1 combination, with the goal of durable tumor control in PTEN/p53-deficient PC.

**Experimental design:** Pb-Cre;PTEN^fl/fl^Trp53^fl/fl^ GEM were treated with either ADT (degarelix), PI3Ki (copanlisib), aPD-1, MEK inhibitor (trametinib) or Porcupine inhibitor (LGK 974) as single agents or their combinations. MRI was used to monitor tumor kinetics and immune/proteomic profiling/*ex vivo* co-culture mechanistic studies were performed on prostate tumors or established GEM-derived cell lines.

**Results:** We tested whether Wnt/β-catenin pathway inhibition with LGK 974 addition to degarelix/copanlisib/aPD-1 therapy enhances tumor control in GEM, and observed *de novo* resistance due to feedback activation of MEK signaling. Based on our observation that degarelix/aPD-1 treatment resulted in partial inhibition of MEK signaling, we substituted trametinib for degarelix/aPD-1 treatment, and observed a durable tumor growth control of PI3Ki/MEKi/PORCNi in 100% mice via H3K18lac suppression and complete TAM activation within TME.

**Conclusions:** Abrogation of lactate-mediated cross-talk between cancer cells and TAM results in durable ADT-independent tumor control in PTEN/p53-deficient AVPC, and warrants further investigation in clinical trials.

**STATEMENT OF TRANSLATIONAL RELEVANCE:** PTEN loss-of-function occurs in ∼50% of mCRPC patients, and associated with poor prognosis, and immune checkpoint inhibitor resistance across multiple malignancies. Our prior studies have demonstrated that ADT/PI3Ki/PD-1 triplet combination therapy controls PTEN/p53-deficient PC in 60% of mice via enhancement of TAM phagocytosis. Here, we discovered that resistance to ADT/PI3K/PD-1 therapy occurred via restoration of lactate production via feedback Wnt/MEK signaling following treatment with PI3Ki, resulting in inhibition of TAM phagocytosis. Critically, co-targeting of PI3K/MEK/Wnt signaling pathways using an intermittent dosing schedule of corresponding targeted agents resulted in complete tumor control and significantly prolonged survival without significant long-term toxicity. Collectively, our findings provide “proof-of-concept” that targeting lactate as a macrophage phagocytic checkpoint controls growth of murine PTEN/p53-deficient PC and warrant further investigation in AVPC clinical trials.

## INTRODUCTION

Prostate cancer (PC) is diagnosed in 283,000 US men annually, with an estimated 34,700 death every year (1). While androgen deprivation therapy (ADT) has remained the cornerstone of therapy for advanced disease, progression to metastatic castrate-resistant prostate cancer (mCRPC) is inevitable. Several FDA approached therapies, including taxane chemotherapy (docetaxel, cabazitaxel), androgen receptor signaling inhibitors (abiraterone, enzalutamide and apalutamide), dendritic cell vaccine (sipuleucel-T) and radiopharmaceuticals (Lutetium-177 vipivotide tetraxetan), have demonstrated modest improvement in overall survival of mCRPC patients (2, 3). However, the disease remains incurable with high morbidity and mortality. Recent intensified approaches with upfront chemohormonal therapies in the metastatic castration-sensitive PC setting have demonstrated survival benefit (2–5). However, it has led to an increased frequency of aggressive-variant forms of the disease in the mCRPC setting, with loss-of-function (LOF) alterations in transformation-related protein 53 (TP53), retinoblastoma (RB) phosphatase and tensin homolog (PTEN) genes, that can drive lineage plasticity and neuroendocrine/small cell histopathological features (6, 7). The development of definitive medicines to treat these aggressive-variant prostate cancers (AVPC) represents an area of critical unmet need.

PTEN LOF genetic alteration occurs in ∼50% of mCRPC patients, which contributes to poor prognosis, therapeutic outcomes and resistance to immune checkpoint inhibitors (ICI) (8–12). Prior studies have demonstrated that PI3K-AKT pathway downstream of PTEN loss stimulates immunosuppressive cytokine release, such as CCL2 and IL-10 (10, 11, 13), resulting in infiltration of regulatory T cells (T_reg_ cells), type-2 helper T cells (Th2 cells), M2-polarized tumor-associated macrophages (M2-TAM), myeloid-derived suppressive cells (MDSC) (10, 11, 14), and decreased frequency of memory CD8^+^ T cells and type-1 helper T cells (Th1 cells) within TME (10, 14). Furthermore, PTEN loss induces expression of programmed death-ligand 1 (PD-L1) and indoleamine 2, 3-dioxygenase 1 (IDO1), which suppress anti-tumor T cell responses within TME (11, 14). Collectively, these PTEN loss-induced immunosuppressive mechanisms contribute to the lack of responsiveness to ICI efficacy across multiple cancers.

Our recent studies have demonstrated that PI3K inhibitor (PI3Ki) suppresses lactate production from PTEN/p53-deficient PC cells, resulting in restoration of TAM phagocytosis and partial suppression of Pb-Cre;PTEN^fl/fl^Trp53^fl/fl^ GEM tumor growth (15), which was enhanced by the addition of ADT/PD-1 blockade. However, upregulation of Wnt/β-catenin signaling following ADT/PI3Ki/aPD-1 therapy led to resistance in 40% of mice via restoration of lactate production from PC cells and corresponding histone lactylation and immunosuppression within TAM (15). In our current study, we hypothesized that suppression of Wnt/β-catenin signaling in combination with ADT/PI3Ki/aPD-1 therapy would overcome lactate-mediated TAM immunosuppression and induce durable PTEN/p53-deficient PC growth control. Following the addition of PORCNi to ADT/PI3Ki/aPD-1therapy in prostate-specific PTEN/p53-deficient GEM, we observed feedback upregulation of MEK signaling, resulting in restoration of lactate production, increased histone lactylation and suppression of TAM-mediated anti-tumor phagocytic response. Given our observation that degarelix/aPD-1 treatment resulted in partial inhibition of MEK signaling, which was insufficient to suppress histone lactylation within TME in the context of ADT/PI3Ki/aPD-1/PORCNi treatment, we substituted a MEK inhibitor for degarelix/aPD-1 treatment, and observed tumor growth control in 100% of mice treated with PI3Ki/MEKi/PORCNi, which was completely abolished by *in vivo* clordronate-mediated depletion of systemic activated TAM. Mechanistically, inhibition of lactate production from PI3Ki/MEKi/PORCNi-treated PC cells suppressed histone lactylation within TAM, resulting in. their phagocytic activation and uptake of PC cells. Given concerns for toxicity following long-term treatment with PI3Ki/MEKi/PORCNi, we developed an intermittent dosing strategy for PI3Ki/MEKi/PORCNi combination treatment, which demonstrated long-term tumor control and favorable safety profile in 100% mice. Collectively, these data warrant further investigation of PI3Ki/MEKi/PORCNi intermittent dosing strategies to activate TAM-mediated innate immunity and produce durable tumor control in PTEN-deficient mCRPC patients.

## MATERIALS AND METHODS

### *In vivo* murine treatment and prostate tumor growth kinetic studies

All studies performed in mice were approved by the Institutional Animal Care and Use Committee (IACUC) at University of Chicago and performed in compliance with NIH guidelines. Prostate-specific PTEN/p53-deficient (Pb-Cre;PTEN^fl/fl^Trp53^fl/fl^) mice were screened for autochthonous prostate tumor development at 16 weeks of age by ultrasound. Following the development of solid tumors (5 mm long-axis diameter of solid tumor with ultrasound imaging), the mice were treated with either degarelix (0.625 mg/mouse, *sc*, every 28-days, ADT group, MedchemExpress; HY16168A), copanlisib (14mg/kg, *iv*, every alternate day, Selleckchem; BAY 80-6946), PD-1 antibody (aPD-1, 200 μg/mouse, *ip*, every alternate day, Bristol Myers Squibb; mPD-1-D265A), trametinib (3mg/kg, *po*, every alternate day, Selleckchem; GSK 1120212), LGK 974 (3mg/kg, *po*, every alternate day, Selleckchem; WNT 974) single agents or their combinations (as indicated in Figure legends). Tumor volumes were monitored using MRI and quantified using Amira software (RRID:SCR_007353), as previously described (15). If solid tumor volume did not grow >20% (stable disease, SD) at indicated time point following treatment, relative to baseline, mice were considered treatment responsive. Percent Partial response (PR) was calculated based on a proportion of total mice that exhibited >30% decrease in solid tumor volume following treatment, relative to baseline. % Overall Response Rate (ORR) was calculated based on number of responder mice (SD + PR), relative to total mice enrolled in a specific treatment.

### Cell lines and culture conditions

AC1 (adenocarcinoma type) and SC1 (sarcomatoid type) cancer cells were derived from PTEN/p53-deficient prostate GEMM tumor and previously authenticated (16). AC1 and SC1 cells were cultured in prostate epithelial growth media (PrEGM), which was generated by adding necessary supplements (Lonza; CC-3166) to Prostate Epithelial Basal Media (PrEBM, Lonza; CC-4177), as per manufacturer’s protocol, in the absence and presence of 10% fetal bovine serum (FBS, Gemini; 100500), respectively (16). These cells were confirmed to be mycoplasma free using PCR-based testing kit (ATCC; 30-1012K).

### Western blot analysis

Established prostate tumors from Pb-Cre;PTEN^fl/fl^Trp53^fl/fl^ mice were harvested following treatment at timespoints indicated in Figure legends, lysed in T-PER buffer (ThermoFisher; 78510) and 10 μg of total protein extracts for each sample underwent sodium dodecyl sulfate-polyacrylamide gel electrophoresis (SDS-PAGE), followed by Western Blot analysis. AC1 or SC1 cells were treated with indicated drug(s), lysed in RIPA buffer (ThermoFisher; 89900) and 10 μg of total protein extracts underwent SDS-PAGE followed by Western blotting, and probed for pAKT-S473 (Cell Signaling Technology; 4060S), pAKT-T308 (Cell Signaling Technology; 2965S), total AKT (Cell Signaling Technology; 4691S), active-β-catenin (Cell Signaling Technology; 8814S), H3K18lac (PTM biolabs; PTM-1406RM) pERK-T202/Y204 (Cell Signaling Technology; 9101S), total ERK (Cell Signaling Technology; 4691S) or GAPDH (Cell Signaling Technology; 4695S) as indicated in Figure legends.

### Flow cytometric analysis

Pb-Cre;PTEN^fl/fl^Trp53^fl/fl^ mice were treated with the indicated drug(s). Single cell suspensions were prepared from treated tumors using liberase (Sigma; 5401020001) as previously described (15), and stained with either myeloid or lymphoid antibody cocktails at a density of 1x10^6^ cells/mL in PBS. Briefly, myeloid antibody cocktails were prepared by resuspending 10 μL of anti-CD45 (Biolegend; 103138), CD11b (Biolegend; 101257), CD11c (Biolegend; 117339), Ly6c (Biolegend; 128006), Ly6g (Tonbo Biosciences; 80-5931-U100), MHC-II (Biolegend; 107631), F4/80 (Biolegend; 123147), CD206 (Biolegend; 141708), PD-1 (Biolegend; 135228), PD-L1 antibodies (Invitrogen; 25-5982-82) and 1 μL of Ghost-viability dye (Tonbo Biosciences; 13-0870-T100) in 1 mL PBS. Lymphoid antibody cocktails were prepared by resuspending 10 μL of anti-CD45, CD3 (Biolegend; 100216), CD4 (Biolegend; 100451), CD8 (Biolegend; 100748), CD19 (Biolegend; 115541), PD-1, PD-L1 antibodies and 1 μL of Ghost-viability dye in 1 mL PBS. Following 30 minutes of surface staining, single cell suspensions were washed with PBS and incubated with fix/perm buffer (Biolegend; 421401) for 15 minutes. After fixation, single cell suspensions of samples incubated with myeloid antibody cocktail were washed and resuspended in PBS for flow cytometry run whereas lymphoid antibody cocktail-stained cells were incubated overnight with 1 mL perm buffer (Biolegend; 421402) containing 10 μL of anti-FoxP3 (Biolegend; 126408) and Ki67 (Biolegend; 652404) antibodies for intracellular staining. Following completion of staining, cells were washed, resuspended in PBS and flow cytometry was performed for lymphoid markers. Flow cytometry data were gated for myeloid and lymphoid subsets and their activation status using FlowJo v 10.7 software (RRID:SCR_008520).

### CM collection

As described above, single cell suspensions were prepared using liberase digestion method using established prostate tumors from untreated Pb-Cre;PTEN^fl/fl^Trp53^fl/fl^ mice, and were plated at density of 3x10^6^ cells per P100-culture dish. These suspensions were treated *ex vivo* with copanlisib (100 nM), trametinib (5 nM), LGK 974 (50 nM) or their combinations under AC1 media conditions, as indicated in Figure legends. Following treatment, supernatants were collected, named as “*ex vivo* CM” and utilized for TAM functionality experiments. For *in vitro* CM, supernatants were collected following treatment of AC1/SC1 cells with copanlisib (100 nM), trametinib (5 nM), LGK 974 (50 nM) or their combinations in presence of AC1/SC1 media, as described in Figure legends.

### Lactate determination

Lactate was quantified using colorimetry kit (Biovision; K627-100) on both *ex vivo* and *in vitro* CM.

### *Ex vivo* phagocytosis assay

TAM or their subsets were sorted from untreated PTEN/p53-deficient established prostate tumors using FACS and utilized for two distinct phagocytosis assay approaches. (i) TAM were either <u>pre-treated</u> directly with copanlisib (100 nM), trametinib (5 nM), LGK 974 (50 nM) or their combinations in presence of normal AC1/SC1 media or (ii) <u>indirectly with CM</u> (as described in above subsection) for 24 hours to dissect mechanism of TAM activation/phagocytosis. To investigate role of lactate on macrophage suppression, FACS-sorted TAM or BMDM were pre-treated for 24 hours with *ex vivo* or *in vitro* CM, in the presence or absence of lactate (Sigma; L7022), which was added at a final concentration of either 50 or 100 nmol/μL After these pre-treatments, TAM were co-cultured with CTV dye stained AC1/SC1 cells at Tumor cells:TAM ratio of 2:1 for 2 hours. At the end of phagocytosis, cells were then washed twice with PBS and stained with anti-CD45, MHC-II (M1-anti-tumor/activated macrophage marker), PD-1, (1:100 dilution for each), H3K18lac antibodies (1:100 dilution for each) and Ghost-viability dye (1:1000 dilution) in 1 mL PBS. Histone lactylation status and phagocytic activity and of each TAM subset (MHC-II^hi^/PD-1^lo^, MHC-II^hi^/PD-1^hi^, MHC-II^lo^/PD-1^lo^, and MHC-II^lo^/PD-1^hi^ TAM) were calculated by normalizing H3K18lac staining and MFI of CTV dye, respectively, relative to untreated groups,. Baseline phagocytic activity of TAM subsets was normalized relative to untreated MHC-II^lo^PD-1^hi^ TAM.

### Proliferation and apoptosis assays

AC1/SC1 cells were treated with copanlisib (100 nM), trametinib (5 nM), LGK 974 (50 nM) or their combinations in presence of AC1/SC1 for indicated time points. Following treatment, proliferation rate was calculated by dividing the number of cells at a given time point with the seeding density. Hemocytometer method was utilized to count cells at any given time point. Furthermore, cells were stained with anti-Annexin V antibodies and propidium iodide (PI, DNA marker dye) as per protocol supplied with kit (BD biosciences; 556547). Flow cytometry analysis was performed for annexin V+/PI-(apoptotic) and annexin V+/PI+ (necrotic) cells. % cell death was defined as the sum total of both apoptotic and necrotic cells frequencies.

### Histopathological and survival analysis

Power calculation was done using published data of Pb-Cre;PTEN^fl/fl^Trp53^fl/fl^ mice (n = 32, mortality = 100% and median survival time = 5.5 months) relative to wild-type mice (n=33, mortality = 0% with 10 months follow-up), with hazard ratio = 0.03099 (17). Based on these estimates, the required number of mice using α = 0.05 and β = 0.2 was n=3 in treated group with no mortality to 11 months, relative to n=3 untreated mice. Pb-Cre;PTEN^fl/fl^Trp53^fl/fl^ mice with established prostate tumors were treated with copanlisib (14 mg/kg, *iv*, every alternate day) + trametinib (3 mg/kg, *po*, every day) + LGK 974 (3 mg/kg, *po*, every day) until signs of distress post-9 weeks of treatment, such as mild hunched, incision, rough hair coat, squinted eye, sudden body weight loss appeared (18). After mice recovered from visible signs of distress following 3 weeks of drug holiday, treatment schedule was resumed again until mice reached 11 months of age, relative to untreated historic controls (100% mortality by 7 months) (17). Tumors were harvested and approximately 5-μm thick formalin-fixed-paraffin-embedded sections were prepared. Hematoxylin and Eosin (H&E) staining was performed on these sections, and histopathological assessments were carried out by a blinded pathologist for anti-tumor efficacy.

## RESULTS

### Prostate tumors in Pb-Cre;PTEN^fl/fl^Trp53fl^/fl^ mice are resistant to ADT/PI3Ki/aPD-1/PORCNi combination therapy due to feedback activation of MEK signaling

Our recent studies have demonstrated Wnt/β-catenin pathway activation in 40% of Pb-Cre;PTEN^fl/fl^Trp53^fl/fl^ mice that have acquired resistance to ADT/PI3Ki/aPD-1 treatment, resulting in restoration of tumor cell intrinsic lactate production and corresponding TAM histone lactylation/phagocytosis suppression (15). As a first step towards understanding whether inhibition of Wnt/β-catenin signaling overcomes this treatment-resistance and restores tumor control in Pb-Cre;PTEN^fl/fl^Trp53^fl/fl^ mice, we tested the therapeutic impact of ADT/PI3Ki/aPD-1 in combination with a porcupine inhibitor, which suppresses Wnt/β-catenin signaling (19). Established tumors from Pb-Cre;PTEN^fl/fl^Trp53^fl/fl^ mice were first treated with ADT/copanlisib (PI3Ki)/aPD-1 combination and monitored for the emergence of resistance by MRI, at which time LGK 974 (Porcupine inhibitor) was added. We observed that LGK’974 was unable to suppress tumor growth following the development of resistance to ADT/copanlisib/aPD-1 combination therapy **(Fig. 1A)**. Western blot analysis revealed feedback upregulation of MEK signaling and an increase in histone lactylation within tumors that were treated with LGK 974 following resistance to ADT/copanlisib/aPD-1 combination therapy **(Fig. 1B)**. Flow cytometry analysis revealed a decrease in the frequency of activated TAM and T-cells within the prostate tumors of ADT/copanlisib/aPD-1/LGK 974 combination treated mice, relative to ADT/copanlisib/aPD-1 responder mice, and no change relative to untreated controls **(Fig. 1C)**. These findings demonstrate that Wnt/β-catenin inhibition was unable to rescue the subset of Pb-Cre;PTEN^fl/fl^Trp53^fl/fl^ mice that are resistant to ADT/PI3Ki/aPD-1 combination therapy, due to feedback upregulation of MEK signaling.

**Figure 1.**
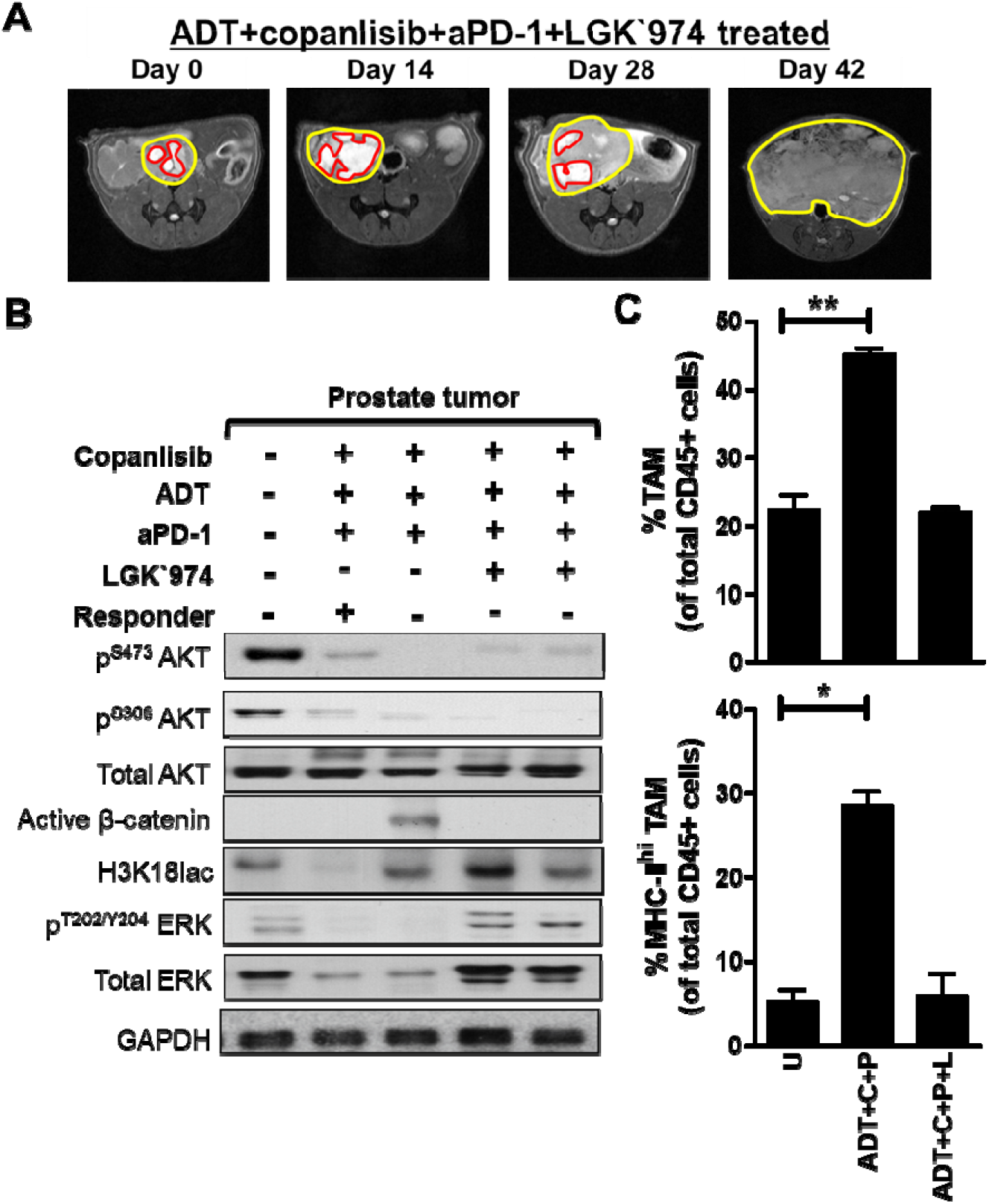
The addition of PORCNi is unable to control tumor growth in Pb-Cre;PTEN^fl/fl^ Trp53^fl/fl^ mice that are resistant to ADT/PI3Ki/aPD-1 combination therapy, due to feedback upregulation of MEK signaling. (A) Pb-Cre;PTEN^fl/fl^ Trp53^fl/fl^ mice with established solid prostate tumors were treated with copanlisib (14 mg/kg, *iv*, every alternate day, 42 days) + degarelix (0.625 mg, every month, 42 days, ADT) + PD-1 antibody (100 μg/dose, every alternate day, 42 days, aPD-1). On day 28, non-responder mice received LGK 974 (porcupine inhibitor, 3 mg/kg, *po*, daily, 14 days). The change in solid tumor volume is qualitatively represented by MRI for ADT + copanlisib+ aPD-1 + LGK 974 treated group. Total tumor area and cystic parts of tumors are outlined by yellow and red colors on MRI, respectively. (B) At the end of 42 days of treatment, tumor extracts obtained from mice treated with indicated drug(s) were profiled by western blot analysis for PI3K, MEK and Wnt signaling. (C) Flow cytometry was performed to assess infiltration frequency (upper panel) and activation status (MHC-II^hi^, lower panel) of tumor-associated macrophages (TAM, CD11b+Ly6g-Ly6c-F4/80+), relative to control mice that responded to ADT + copanlisib + aPD-1 treatment. Bar graph represents. n=2 mice per group. Significances/p-values were calculated by one-way ANOVA and indicated as follows, *p<0.05 and **p<0.01.

### ADT/aPD-1 combination inhibits MEK signaling in PTEN/p53-deficient prostate tumors

In addition to inhibition of PI3K signaling, we observed that ADT/PI3Ki/aPD-1 combination suppressed MEK signaling within PTEN/p53-deficient prostate tumors, relative to untreated control **(Fig. 1B)**. In contrast, MEK pathway activation emerged as a resistance mechanism to ADT/PI3Ki/aPD-1/LGK 974 combination treatment, but not with ADT/PI3Ki/aPD-1 combination **(Fig. 1B)**. To elucidate this paradox, we first tested the hypothesis that ADT/aPD-1 combination therapy inhibits MEK pathway activation within TME of Pb-Cre;PTEN^fl/fl^Trp53^fl/fl^ mice. We performed Western blot analysis on prostate tumors derived from ADT/aPD-1 combination treated mice, which showed decreased p-ERK, relative to untreated control, indicative of suppression of MEK signaling. In contrast, there was no significant change in p-AKT or active β-catenin with the combination, markers of PI3K and Wnt/β-catenin signaling respectively, relative to untreated controls **(Fig 2A)**. Collectively, these data demonstrate that dual ADT/aPD-1 treatment suppresses MEK signaling within PTEN/p53-deficient GEMM tumors, which is reversed by addition of LGK 974 in combination with ADT/PI3Ki/aPD-1 therapy.

**Figure 2.**
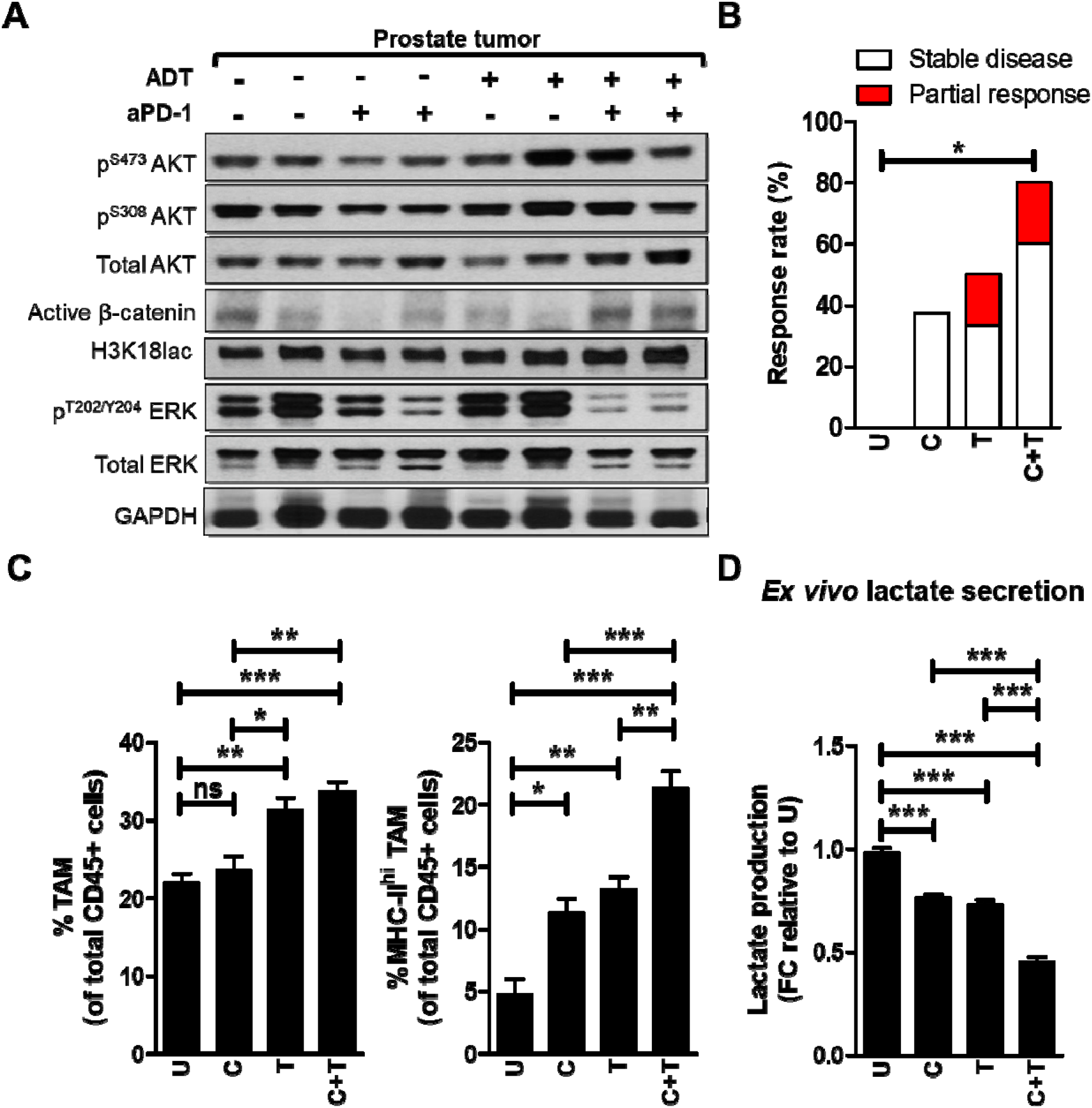
PI3Ki/MEKi combination therapy induces activated TAM-mediated tumor growth control in 80% of Pb-Cre;PTEN^fl/fl^Trp53^fl/fl^ mice. (A) Pb-Cre;PTEN^fl/fl^Trp53^fl/fl^ mice with established prostate tumors were treated with PD-1 antibody (aPD-1, 100 μg/mouse, *ip*, every alternate day) alone or in combination with ADT (degarelix, 0.625mg, single dose, castration) for 28 days. Western blot analyses were performed for indicated protein markers on prostate tumor extracts. (B) Pb-Cre;PTEN^fl/fl^Trp53^fl/fl^ mice with established prostate tumors were treated with trametinib (3 mg/kg, *po*, every day), singly and in combination with copanlisib (14 mg/kg, *iv*, every alternate day) for 28 days. Tumor growth was monitored using MRI and response rates (Partial response plus Stable disease) were quantified as described in Methods. (C) At the end of treatment, prostate tumors were profiled using flow cytometry and analyzed for % frequency of total and MHC-II expressing TAM (CD45^+^CD11b^+^F4/80^+^ cells). (D) Single cell suspensions of PTEN/p53-deficient prostate GEM tumors were treated with copanlisib (C, 100 nM), trametinib (T, 5 nM) or their combination for 24 hours as described in Methods. *Ex vivo* CM were collected from these groups and analyzed for lactate content using colorimetry kits. For *in vivo* studies, n=5-6 mice per group and for *ex vivo*, n=2 independent experiments. Significances/p-values were calculated by Chi-square test (panel B), one-way ANOVA (panel C-D) and indicated as follows, *p<0.05, **p<0.01 and ***p<0.001; ns = not statistically significant.

### Combination of PI3K and MEK inhibitors control tumor growth in 80% of Pb-Cre; PTEN^fl/fl^ Trp53^fl/fl^ mice via suppression of histone lactylation within TAM

Based on our observation that ADT/aPD-1 combination partially suppresses MEK signaling within tumors *in vivo*, we tested the hypothesis that MEKi (substituted for ADT/aPD-1 in the ADT/PI3Ki/aPD-1 backbone) will lead to at least equivalent tumor control via TAM activation. Due to toxicity reported with PI3Ki/MEKi combination in clinical trials (20), we first evaluated the safety and efficacy of copanlisib and trametinib (MEKi) combination treatment in Pb-Cre;PTEN^fl/fl^Trp53^fl/fl^ mice with established prostate tumors for 4 weeks. Trametinib alone, which completely suppressed intratumoral MEK signaling led to a 50% response rate, relative to the 33.3% response rate observed with ADT/aPD-1 doublet therapy. Importantly, the addition of copanlisib to trametinib increased response rate to 80% **(Fig. 2B and Supplementary Fig. S1A-B),** relative to the 60% response rate observed with the ADT/PI3Ki/aPD-1 combination, with equivalent partial response rates (15). Immune profiling demonstrated a 2.8-fold increase in frequency of activated TAM within prostate tumors from trametinib-treated mice, which increased to a 4.6-fold with trametinib/copanlisib, relative to untreated control **(Fig. 2C)**.

We have previously demonstrated that ADT/PI3Ki/aPD-1 combination controls tumor growth in Pb-Cre;PTEN^fl/fl^Trp53^fl/fl^ mice via suppression of histone lactylation and corresponding enhancement of TAM phagocytosis, particularly in MHC-II^hi^/PD-1^hi/lo^ TAM subsets (15). To determine whether PI3Ki/MEKi combination-induced tumor control is mediated via TAM innate immune phagocytic mechanism, we analyzed *ex vivo* CM collected from single cell suspensions of tumors following treatment with trametinib alone at IC_90_ concentrations **(Supplementary Fig. S2A-B)**, which showed a 24.7% decrease in lactate levels that was further decreased to 72% in combination with copanlisib **(Fig. 2D)**. Importantly, the combination of copanlisib and trametinib demonstrated a 17.2 and 21-fold suppression of histone lactylation in MHC-II^hi^/PD-1^lo^ TAM and MHC-II^hi^/PD-1^hi^ TAM, respectively, relative to untreated. In contrast, single-agent trametinib or copanlisib led to 3.9- and 4-fold suppression of histone lactylation in MHC-II^hi^/PD-1^lo^ and MHC-II^hi^/PD-1^hi^ TAM, respectively, relative to untreated **(Fig. 3A-B and Supplementary Fig. S3A)**, Critically, the combination of copanlisib and trametinib demonstrated a 7.2- and 7.1-fold increase in phagocytic capacity of MHC-II^hi^/PD-1^lo^ and MHC-II^hi^/PD-1^hi^ TAM, respectively, relative to untreated controls. In contrast, single agent trametinib or copanlisib led to a 3.8- and 3.7-fold induction in phagocytic capacity of MHC-II^hi^/PD-1^lo^ and MHC-II^hi^/PD-1^hi^ TAM, respectively, relative to untreated controls **(Fig. 3C and Supplementary Fig. S3B)**. Collectively, these data demonstrate that the PI3Ki/MEKi combination potently suppresses histone lactylation and enhances phagocytosis within activated TAM, relative to untreated or single agent controls. In contrast, direct treatment with trametinib did not alter TAM phagocytic capacity **(Supplementary Fig. S4A-B)**. Furthermore, *in vitro* treatment with copanlisib + trametinib modestly decreased proliferation and survival of AC1 cells, but not SC1 cells **(Supplementary Fig. S5A-B)**, thus demonstrating that the majority of the anti-cancer response observed is mediated via tumor cell extrinsic immune mechanism(s).

**Figure 3.**
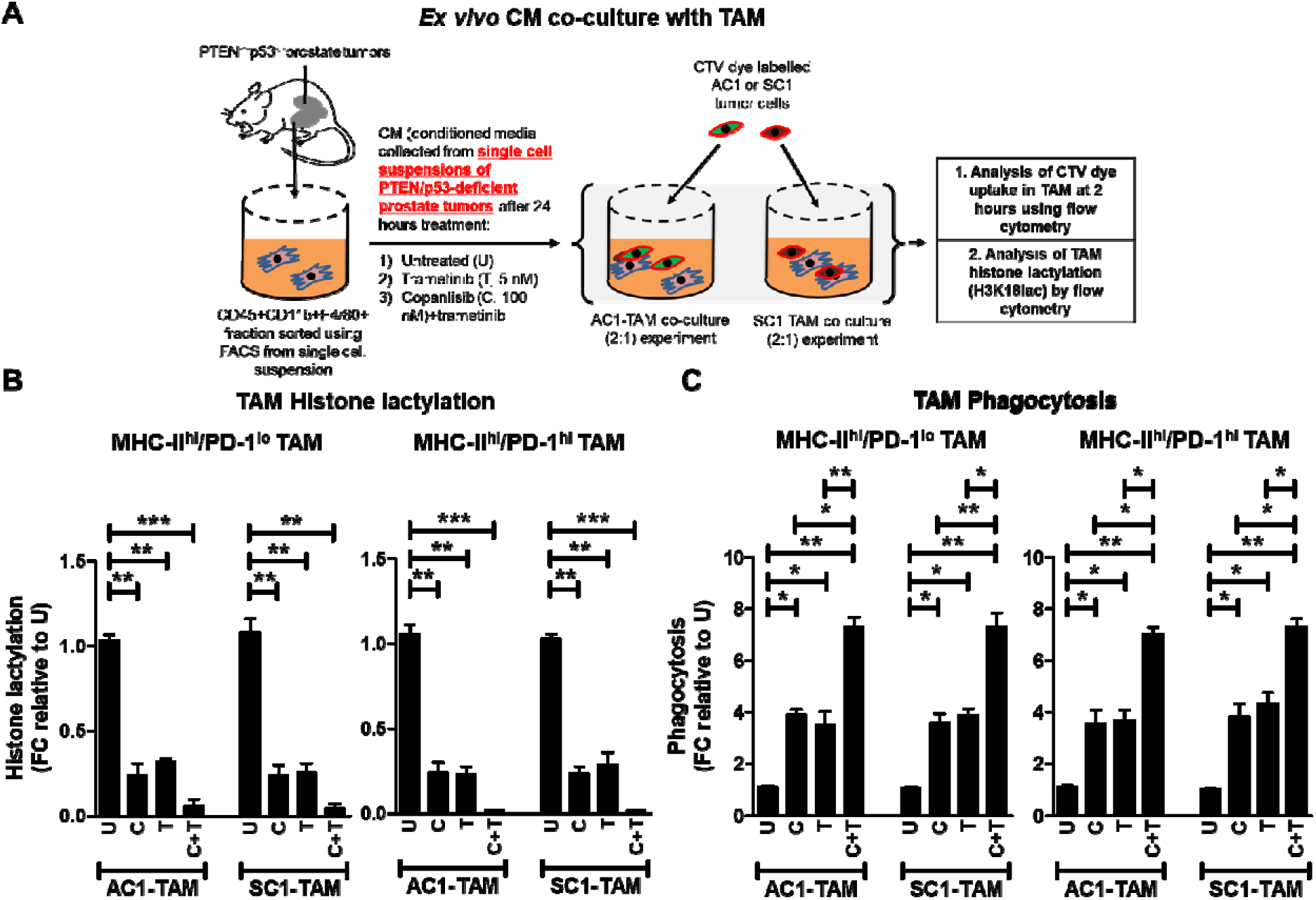
PI3Ki/MEKi combination therapy potently suppresses histone lactylation and enhances phagocytosis within activated TAM, relative to single agent controls. (A) Single cell suspensions of PTEN/p53-deficient prostate GEM tumors were treated with copanlisib (C, 100 nM), trametinib (T, 5 nM) or their combination for 24 hours, and conditioned media (CM) was collected at the end of treatment. FACS-sorted TAM were incubated *ex vivo* in CM for 24 hours, followed by co-culture with CTV dye stained-AC1/SC1 cells for 2 hours. Bar graphs demonstrate histone lactylation status (B) and phagocytic activity (C) of MHC-II^hi^/PD-1^hi/lo^ expressing TAM, relative to untreated group. FC = fold change. n=2 independent experiments. Significances/p-values were calculated by one-way ANOVA and indicated as follows, *p<0.05, **p<0.01 and ***p<0.001.

### Feedback activation of Wnt/**β**-catenin signaling observed in non-responder Pb-Cre;PTEN^fl/fl^Trp53^fl/fl^ mice following PI3Ki/MEKi combination treatment, restores histone lactylation (H3K18lac) and suppresses macrophage phagocytosis

To assess the mechanism of resistance in 20% of mice treated with PI3Ki/MEKi combination therapy, Western blot analysis was performed on whole tumor extracts, which demonstrated an increase in active β-catenin expression and histone lactylation within tumor from non-responder outlier mouse, relative to responder and untreated groups **(Fig. 4A)**. Consistent with these observations, *in vitro* western blot analysis of PTEN/p53-deficient tumor-derived AC1/SC1 cells following 72-hours treatment of copanlisib/trametinib combination demonstrated a concomitant increase in active β-catenin expression, relative to 24-hours treatment **(Supplementary Fig. S6A-B)**. This feedback activation of Wnt/β-catenin signaling was accompanied by an increase in lactate production at 72 hours, following an initial decrease with acute copanlisib/trametinib treatment *in vitro* **(Fig. 4B)**. Collectively, these data suggest that resistance to PI3Ki/MEKi combination therapy is driven by upregulation of Wnt-β-catenin signaling and restoration of tumor cell lactate production/TAM histone lactylation.

**Figure 4.**
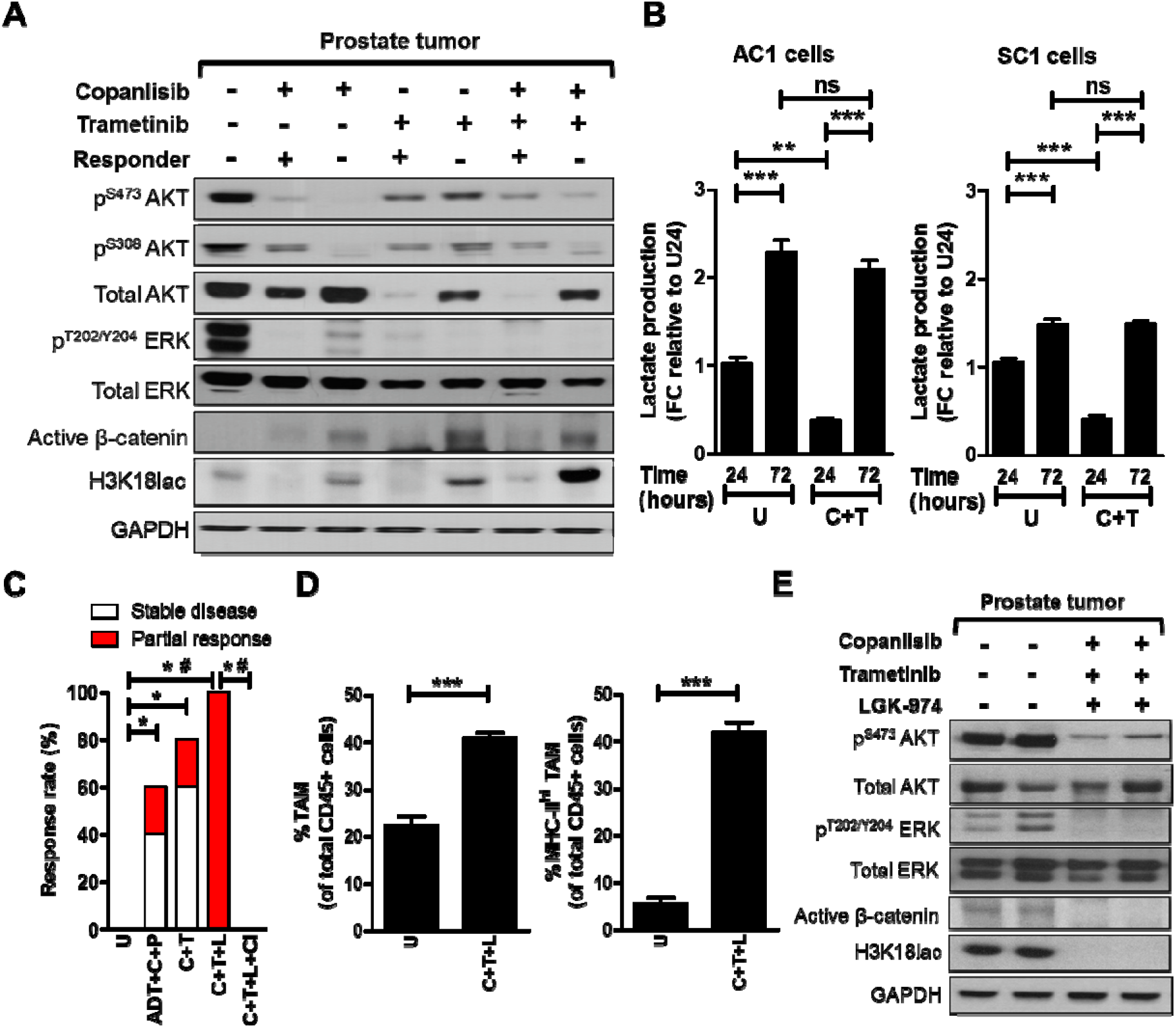
Addition of PORCNi overcomes resistance to PI3Ki/MEKi combination therapy and induces activated TAM-mediated 100% response rate in Pb-Cre;PTEN^fl/fl^Trp53^fl/fl^ mice. (A) Pb-Cre;PTEN^fl/fl^Trp53^fl/fl^ mice with established prostate tumors were treated with trametinib (3 mg/kg, *po*, every day), singly and in combination with copanlisib (14 mg/kg, *iv*, every alternate day) for 28 days. Western blot analyses were performed for indicated protein markers on PTEN/p53-deficient prostate tumor extracts. (B) PTEN/p53-deficient GEM tumor-derived AC1/SC1 cells were treated with copanlisib (C, 100 nM), trametinib (T, 5 nM) or their combination for 24 and 72 hours. *In vitro* CM were collected following treatment and analyzed for lactate content using colorimetry kits. (C) Pb-Cre;PTEN^fl/fl^Trp53^fl/fl^ mice with established prostate tumors were treated with indicated combinations of trametinib (T, 3 mg/kg, *po*, every day), copanlisib (C, 14 mg/kg, *iv*, every alternate day), LGK 974 (L, 3 mg/kg, *po*, every day), PD-1 antibody (aPD-1, 100μg/mouse, *ip*, every alternate day) and ADT (degarelix, 0.625 mg, single dose) for 28 days. Tumors were monitored with MRI and response rates/partial response/stable disease were determined as described in Methods. Tumors from untreated or copanlisib+trametinib+LGK 974-treated mice were analyzed by flow cytometry for % frequency of total and MHC-II expressing TAM (CD45^+^CD11b^+^F4/80^+^ cells) (D) or analyzed for the indicated proteins by Western blotting (E). n = 5-6 mice per group for in vivo studies; for *in vitro* experiments, n = 3 independent experiments. Significances/p-values were calculated by one-way ANOVA (panel B and D), Chi-square test (panel C) and indicated as follows, *p<0.05, **p<0.01, ***p<0.001 and ^#^p<0.05 for partial response (panel A); ns = not statistically significant.

### PI3Ki + MEKi + PORCNi combination therapy drives tumor shrinkage in 100% of Pb-Cre;PTEN^fl/fl^p53^fl/fl^ mice via complete TAM activation

To overcome Wnt/β-catenin-mediated resistance to PI3Ki/MEKi combination therapy, Pb-Cre;PTEN^fl/fl^Trp53^fl/fl^ mice with established tumors were treated with copanlisib + trametinib + LGK 974, which resulted in a 100% response rate/tumor shrinkage, relative to untreated group **(Fig. 4C and Supplementary Fig. S7A)**. This anti-tumor response was rescued by concurrent clodronate treatment **(Fig. 4C and Supplementary Fig. S7B)** and accompanied by a 2-fold increase in total TAM frequency, relative to untreated. Strikingly, copanlisib + trametinib + LGF’974 resulted in activation of 100% TAM within the TME, relative to approx. 20% activated TAM in untreated tumors **(Fig. 4D)** and a corresponding suppression of histone lactylation within the TME **(Fig. 4E)**. Analysis of *in vitro* CM collected following treatment of AC1 and SC1 cells with copanlisib + trametinib + LGK 974 at pre-determined IC_90_ concentrations **(Supplementary Fig. S2 and Supplementary Fig. S8A-B)** showed a 78% and 50% decrease in lactate levels, respectively **(Supplementary Fig. S9A)**. Furthermore, *in vitro* treatment with copanlisib + trametinib + LGK 974 modestly inhibited the proliferation and survival of PTEN/p53-deficient GEM tumor-derived AC1 cells but not SC1 cells **(Supplementary Fig. S9B-C)**, suggesting a predominantly tumor cell extrinsic mechanism by which the combination exerts its anti-tumor effects *in vivo*. Direct treatment of TAM with copanlisib + trametinib + LGK 974 did not alter TAM phagocytic capacity or histone lactylation status **(Supplementary Fig. S10A-C)**. Critically, co-culture of TAM with CM collected following copanlisib + trametinib + LGK 974 treatment of AC1 and SC1 cells demonstrated a 3.9-fold and 3.1-fold reduction in histone lactylation, with a corresponding 6.8 and 6.3-fold increase in respective MHC-II^hi^/PD-1^lo^ and MHC-II^hi^/PD-1^hi^ TAM phagocytic capacity, respectively, relative to untreated controls. Lactate “add-back” to the CM collected following copanlisib + trametinib + LGK 974 treatment restored histone lactylation and suppressed phagocytic activity of TAM. Furthermore, the addition of LGK’974 to CM following collection from tumor cells treated with copanlisib + trametinib did not result in suppression of histone lactylation or increase in phagocytic activity, relative to untreated, thus demonstrating that tumor cell intrinsic Wnt/β-catenin signaling and lactate production is responsible for histone lactylation and immunosuppression within TAM **(Fig. 5A-B).** In contrast, conditioned media from PI3Ki + MEKi + PORCNi treated PTEN/p53-deficient tumor derived cells does not alter histone lactylation and phagocytic activity of MHC-II^lo^ TAM **(Supplementary Fig. S11A-B)**. Given the toxicity concerns with combinatorial PI3Ki/MEKi/Porcupine inhibitor targeted therapy, we evaluated efficacy and safety of copanlisib + trametinib + LGK 974 with long-term treatment. We observed dermatitis and conjunctivitis in Pb-Cre;PTEN^fl/fl^p53^fl/fl^ mice following 9 weeks of triplet therapy. However, a 3-week drug holiday was sufficient for resolution of dermatitis and conjunctivitis, so we decided to administer the copanlisib + trametinib + LGK 974 regimen on a 9-weeks on/3 weeks off intermittent dosing schedule **(Fig. 6A)**, which significantly improved survival in the Pb-Cre;PTEN^fl/fl^p53^fl/fl^ mice with 100% survival at 11 months (statistically pre-determined endpoint), relative to untreated (100% mortality by 7 months, **Fig. 6B**) (17). Critically, we observed a near-complete pathologic response in responsive mice at the end of both short-term (4 weeks) and long-term (approx. 7 months) treatment **(Fig. 6C and Supplementary Table S1)**. Collectively, these data demonstrate that the decrease in lactate production from PTEN/p53-deficient PC cells in response to PI3Ki + MEKi + Porcupine inhibitor treatment overcomes histone lactylation mediated TAM suppression and stimulates phagocytosis of PC cells by TAM, resulting in long-term tumor control in 100% of the mice, with manageable toxicity in the context of an intermittent dosing schedule **(Fig. 6D)**.

**Figure 5.**
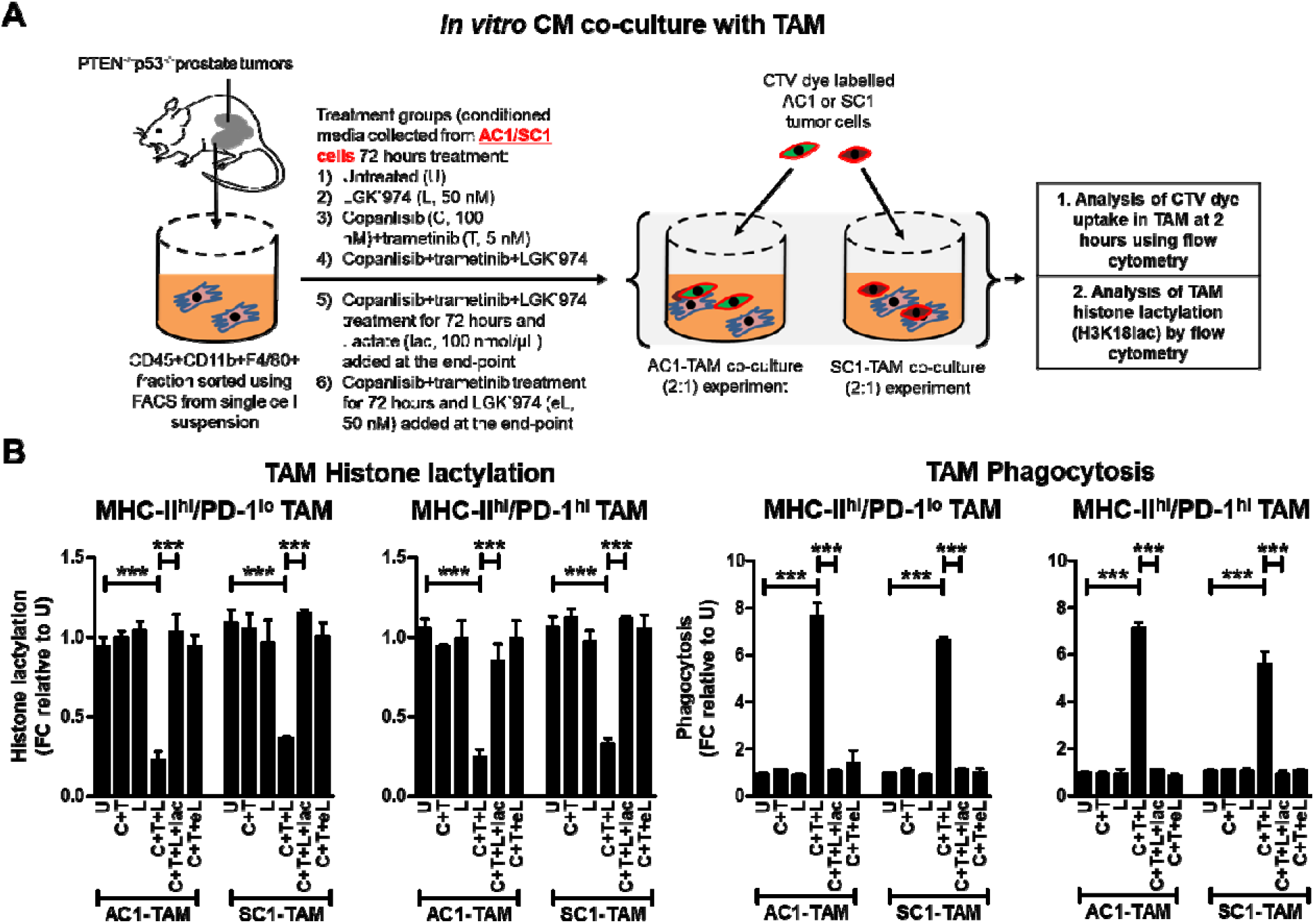
PI3Ki/MEKi/PORCNi combination therapy suppresses lactate production from PTEN/p53-deficient GEM tumor-derived PC cells and secondary histone lactylation within activated TAM, resulting in enhanced TAM phagocytosis. (A) AC1/SC1 cells were treated with copanlisib (C, 100 nM), trametinib (T, 5 nM), LGK 974 (L, 50 nM) or their combination for 72 hours. For mechanistic dissection, lactate (lac, 100 nmol/μL) and LGK 974 (eL, 50nM) were added to the CM collected after C+T+L and C+T treatments of AC1/SC1 cells, respectively. FACS-sorted TAM were incubated with the indicated CM for 24 hours followed by co-culture with CTV dye stained-AC1/SC1 cells for 2 hours. (B) Bar graphs demonstrate fold change (FC) in histone lactylation and phagocytic activity of MHC-II^hi^/PD-1^hi/lo^ expressing TAM, relative to untreated group. n = 2 independent experiments. Significances/p-values were calculated by one-way ANOVA and indicated as follows, ***p<0.001.

**Figure 6.**
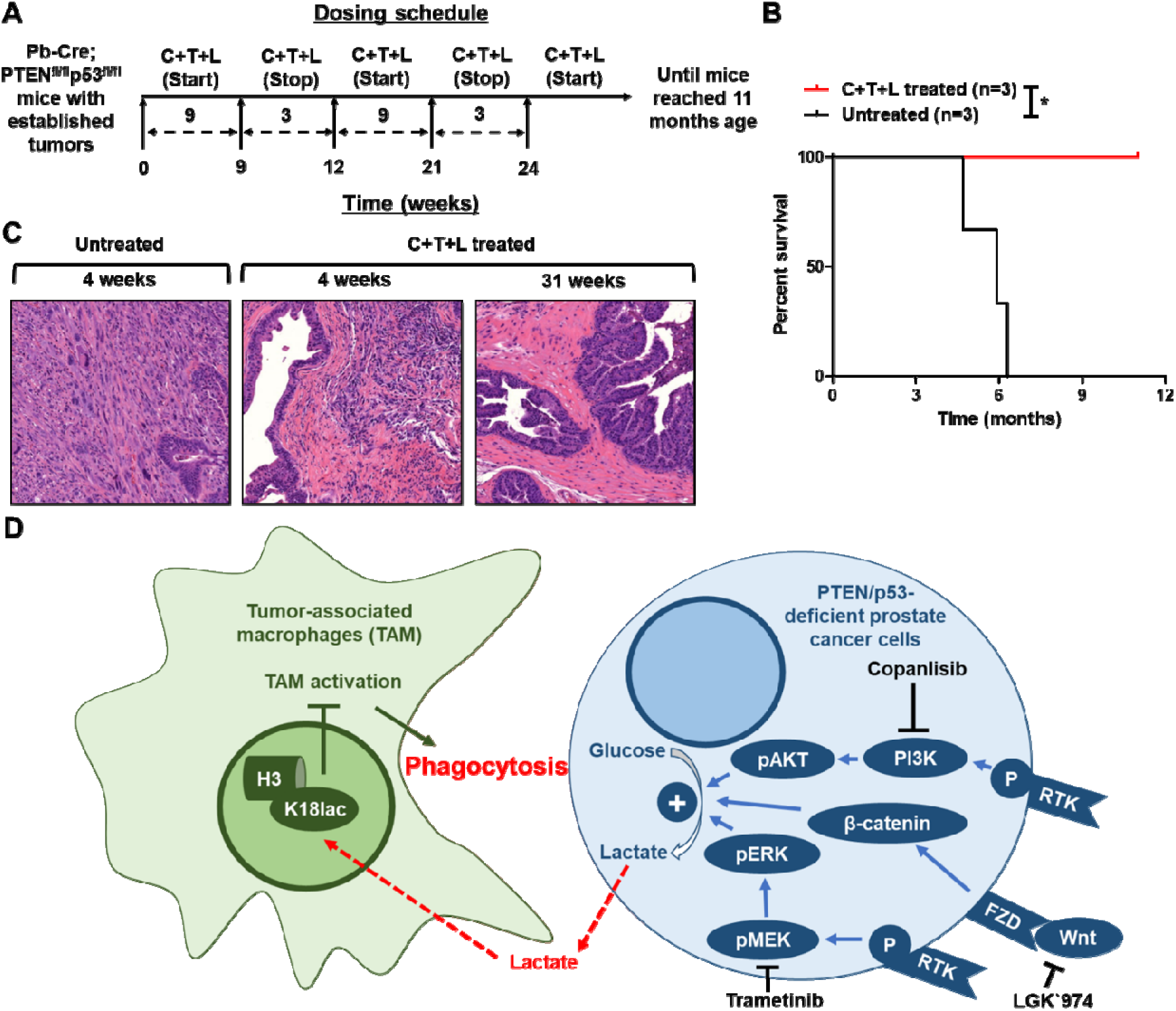
Intermittent long-term dosing of PI3Ki + MEKi + PORCNi results in tumor clearance of established tumors and significant prolongation of survival in Pb-Cre;PTEN^fl/fl^ Trp53^fl/fl^ mice. (A) Schema illustrating Pb-Cre;PTEN^fl/fl^ Trp53^fl/fl^ mice dosed intermittently with copanlisib (C, 14 mg/kg, *iv*, every alternate day) + trametinib (T, 3 mg/kg, *po*, every day) + LGK 974 (L, 3 mg/kg, *po*, every day). (B) Kaplan-Meier survival curves were plotted for C+T+L treatment, relative to untreated control. (C) Tumors were harvested following short-term and long-term C+T+L treatment, and representative histopathologic H&E staining of formalin-fixed paraffin-embedded tissue is depicted. (D) Model illustrating the decrease in lactate production from PTEN/p53-deficient PC cells in response to PI3Ki+MEKi+PORCNi treatment suppresses histone lactylation and enhances phagocytosis of PC cells within activated TAM. n=3 mice per group. Significances/p-values were calculated by Log-rank test and indicated as follows, *p<0.05.

## DISCUSSION

Although ICI have revolutionized the landscape of cancer therapeutics, most mCRPC patients are *de novo* resistant to these medicines, in part due to a paucity of T cells within the TME and/or a tumor cell intrinsic genomic alterations that evade anti-cancer immunity (11, 21). PTEN LOF is present in ∼50% of mCRPC patients that positively correlates with poor prognosis, worse therapeutic outcomes, and ICI resistance (8, 9, 11). Prior studies have demonstrated that hyperactivated PI3K signaling in the context of PTEN LOF TME upregulates GLUT1/4 and hexokinase-2, which enhances aerobic glycolysis. We have previously demonstrated that increased lactate production from PTEN/p53-deficient prostate cancer cells epigenetically reprograms TAM via histone lactylation, and polarizes them towards an immunosuppressive phenotype (M2/MHC-II^lo^ TAM) (15). Furthermore, we identified a negative correlation between TAM M1 activity and tumor cell aerobic glycolysis within the TME of mCRPC patients (15). Metabolically, lactate decreases glycolysis and increases TCA cycle within M1-/M2-TAM, respectively, ultimately reprogramming TAM into a tumor-promoting M2 phenotype (22, 23). Furthermore, lactate augments M2-TAM functionality by upregulating Arg1 expression, resulting in degradation of arginine and decrease in T cell receptor expression and T cell proliferation/effector function (22).

PI3Ki monotherapy has demonstrated limited anti-cancer efficacy in solid tumor malignancies, with or without PTEN/PI3K pathway alterations (24–26). In PTEN-deficient mCRPC patients, co-targeting PI3K and AR signaling with ipatasertib and abiraterone (androgen receptor signaling inhibitor), respectively, demonstrated a modest improvement in progression-free survival (27). Furthermore, our recent preclinical studies in Pb-Cre;PTEN^fl/fl^p53^fl/fl^ mice demonstrated enhanced anti-tumor responses with addition of aPD-1 blockade to the ADT/PI3Ki treatment backbone (15). However, resistance was observed in 40% of mice due to feedback upregulation of Wnt/β-catenin signaling, which restored lactate production and histone lactylation and suppressed phagocytosis within TAM (15). Here, we discovered that PI3K, MEK and Wnt/β-catenin signaling pathways-mediated cross-talk preserves tumor cell-intrinsic lactate production with corresponding TAM histone lactylation/phagocytosis suppression and resultant evasion of anti-cancer innate immunity within PTEN/p53-deficient PC. Concurrent inhibition of these pathways abrogated lactate production from PC cells, resulting in suppression of histone lactylation and phagocytosis-mediated tumor control by activated TAM in 100% of mice. Collectively, we have discovered the importance of lactate as a novel TAM phagocytosis inhibitory checkpoint that evades anti-cancer innate immunity in the context of PTEN/p53-deficient PC. From a translational standpoint, co-targeting oncogenic signaling pathways that preserve lactate production, represents a novel TAM-dependent innate immunomodulatory strategy, that warrants further investigation in combination with ICI in PTEN/p53-deficient AVPC.

Clinically, combination of PI3K and MEK signaling pathway inhibitors have met with challenges related to low therapeutic index and dose-limiting toxicities in clinical trials, including hyperglycemia, loss of appetite, diarrhea, fatigue, rash, vomiting, and increase in lipase and creatine phosphokinase levels (20, 28, 29). A prior preclinical study has demonstrated that transient/intermittent exposure to PI3Ki/MEKi combination is sufficient to induce tumor cell apoptosis relative to continuous administration (30). In this study, sustained pharmacodynamic elevation of Bim protein was found to be critical for apoptosis induction within TME despite recovery of pAKT and pERK following transient exposure to PI3Ki/MEKi combination. These data demonstrated the feasibility of an intermittent administration approach that may be effective to achieve an anti-tumor response while decreasing drug-induced toxicity (30). In our current preclinical *in vivo* studies, an intermittent dosing schedule of PI3Ki/MEKi/Wnti combination (∼9 weeks on and ∼3 weeks off) for at least 7 months led to significant TAM activation/phagocytosis-mediated tumor control in PTEN/p53-deficient murine PC growth, without significant toxicities or mortality. These findings suggest a potential therapeutic window that is achievable with an intermittent dosing approach of PI3Ki/MEKi/PORCNi, which will need further evaluation in Phase I clinical trials. The availability of hyperpolarized ^13^C-pyruvate magnetic resonance spectroscopy imaging approaches would enable *in vivo* pharmacodynamic monitoring of lactate suppression/restoration to optimize intermittent dosing approaches (31). Furthermore, combining imaging with ^13^C-glucose infusion/isotope tracing would provide a safe and effective approach to understand heterogeneity in tumor metabolism, and unravel the immunometabolic mechanism(s) that underly response vs resistance (32).

Alternatively, direct activation of TAM phagocytosis is an optimal strategy to treat PTEN-deficient mCRPC/AVPC patients. This can be achieved by either inhibiting TAM phagocytic checkpoints (15) or blocking lactate entry within TAM (33). In prior published work, we discovered an important role for TAM PD-1 expression in evasion of phagocytosis in Pb-Cre;PTEN^rl/fl^p53^fl/fl^ mice (15). There are several additional TAM checkpoints under investigation, such as SIRPα/CD47 (34), LILRB1/MHC-I (35) and Siglec-10/CD24 (36). Elucidating the functional role of these “don’t eat me” signals in the lactate-rich TME downstream of PTEN loss is an area of unmet need. Furthermore, monocarboxylate transporter (MCT-1, blocks lactate import within TAM) and/or MCT-4 (blocks lactate export from tumor cells) inhibition, which are in early drug discovery and development, could represent promising approaches to directly target lactate-mediated immunometabolic cross-talk within the PTEN-deficient TME of AVPC (33, 37), thereby mitigating potential toxicities encountered with kinase inhibitor cocktails (37).

Clinical trial studies with the porcupine inhibitor, LGK 974, have been limited due to the observed toxicity of increased bone resorption (38, 39). This is of particular concern in mCRPC patients who are on life-long ADT that results in a decreased bone mineral density and increased risk of skeletal-related events (40, 41). Given the complexity of the Wnt/β-catenin signaling pathway, which involves multiple Wnt-ligands, frizzled receptors, co-receptors and extracellular regulators (42), a deeper understanding of Wnt signaling in tumor *vs* bone cells would provide new avenues to effectively treat AVPC patients and mitigate known toxicities. For example, DKK1 blocking agent, which would specifically enhance osteoblast Wnt//β-catenin signaling (43), in combination with PI3Ki/MEKi/PORCNi therapy could be a potential strategy for treating bone resorption in PTEN-deficient mCRPC/AVPC patients. Collectively, our findings provide a mechanistic foundation for a metronomic dose optimization trial to determine immunogenic or TAM activating doses of PI3Ki, MEKi or Wnti agents for treatment of PTEN/p53-deficient AVPC.

## ACKNOWLEDGEMENTS

We thank Drs. Andrew Hsieh, Scott Oakes and Sam Grimaldo for providing helpful suggestions for this manuscript.

## Author s contributions

**K. Chaudagar:** Conceptualization, data curation, formal analysis, validation, investigation, visualization, methodology, supervision, project administration, writing–original draft, writing-review and editing. **H. M. Hieromnimon:** Conceptualization, data curation, formal analysis, validation, investigation, visualization, methodology, writing–original draft, writing-review and editing. **A. Kelley:** Formal analysis, validation, methodology, writing-review and editing. **B. Labadie:** Data curation, visualization, writing–original draft, writing-review and editing. **J. Shafran:** Data curation, visualization, writing–original draft, writing-review and editing. **S. Rameshbabu:** Methodology, writing-review and editing. **C. Drovetsky:** Methodology, writing-review and editing. **K. Bynoe:** Methodology. **A. Solanki:** Methodology. **E. Markiewicz:** Methodology. **M. Zamora:** Methodology. **X. Fan:** Data curation, methodology, writing-review and editing. **M. Loda**: Formal analysis, validation. **A. Patnaik:** Conceptualization, data curation, resources, funding acquisition, supervision, project administration, writing–original draft, writing-review and editing.

