## Supplementary figures for "Suppression of tumor cell lactate-generating signaling pathways eradicates murine PTEN/p53-deficient aggressive-variant prostate cancer via macrophage phagocytosis"

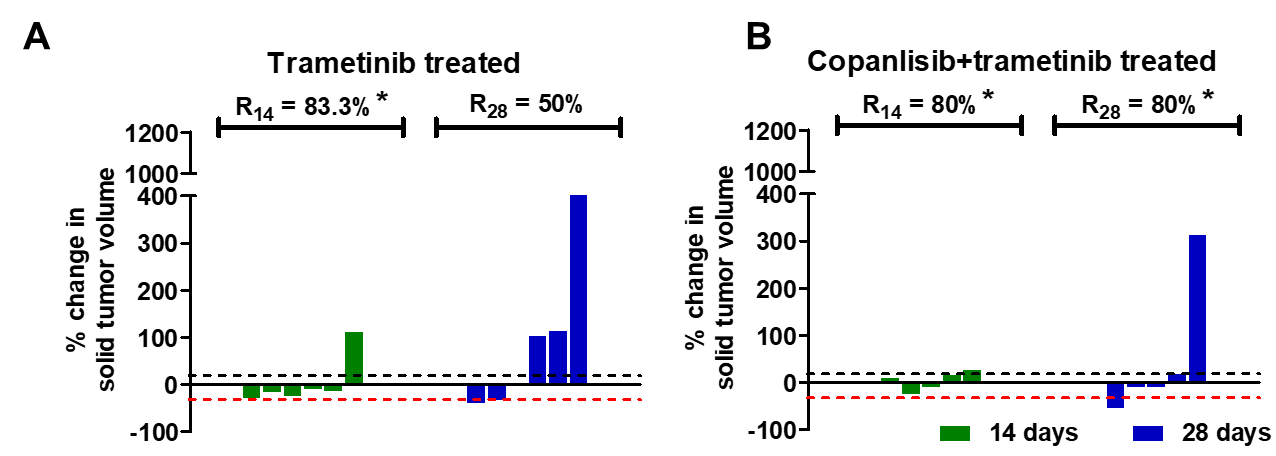

**Supplementary Figure S1. PI3Ki/MEKi combination therapy drives tumor control in majority of Pb-Cre;PTEN^fl/fl^Trp53^fl/fl^ mice.** (A-B) Pb-Cre;PTEN^fl/fl^Trp53^fl/fl^ mice were treated with trametinib (3 mg/kg, *po*, every day) alone and in combination with copanlisib (14 mg/kg, *iv*, every alternate day). Tumor volumes were non-invasively monitored by MRI and % response rate at days 14 (R_14_) and 28 (R_28_) were determined, as described in Methods. The % change in solid tumor volume are represented by waterfall plot trametinib (A) and copanlisib + trametinib (B) treated groups. n=5-6 mice per group. n=5-6 mice per group. Significances/p-values were calculated by Chi-square test and indicated as follows, *p<0.05, (relative to untreated).

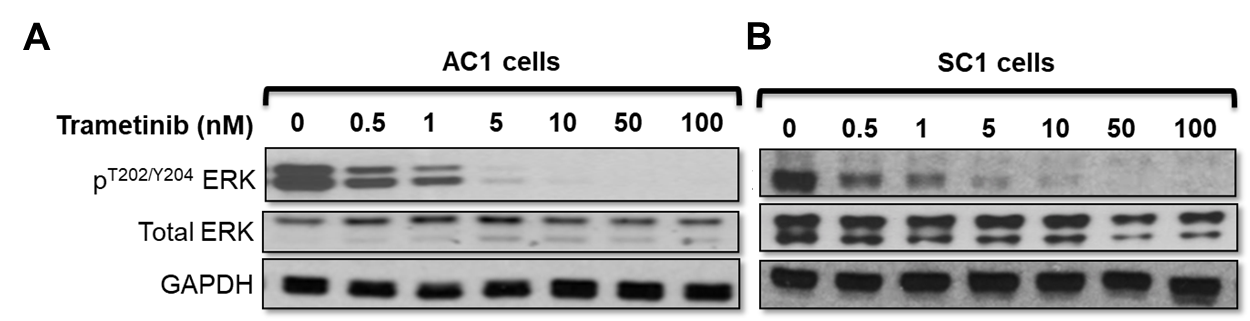

**Supplementary Figure S2. Trametinib inhibits MEK signaling in PTEN/p53-deficient GEM tumor-derived PC cells.** PTEN/p53-deficient GEM tumor-derived PC cells, AC1 (A) and SC1 (B) were treated with trametinib (0-500 nM, 24 hours) in a dose-dependent manner and western blot analyses were performed for indicated proteins to determine IC_90_. n=2 independent experiments.

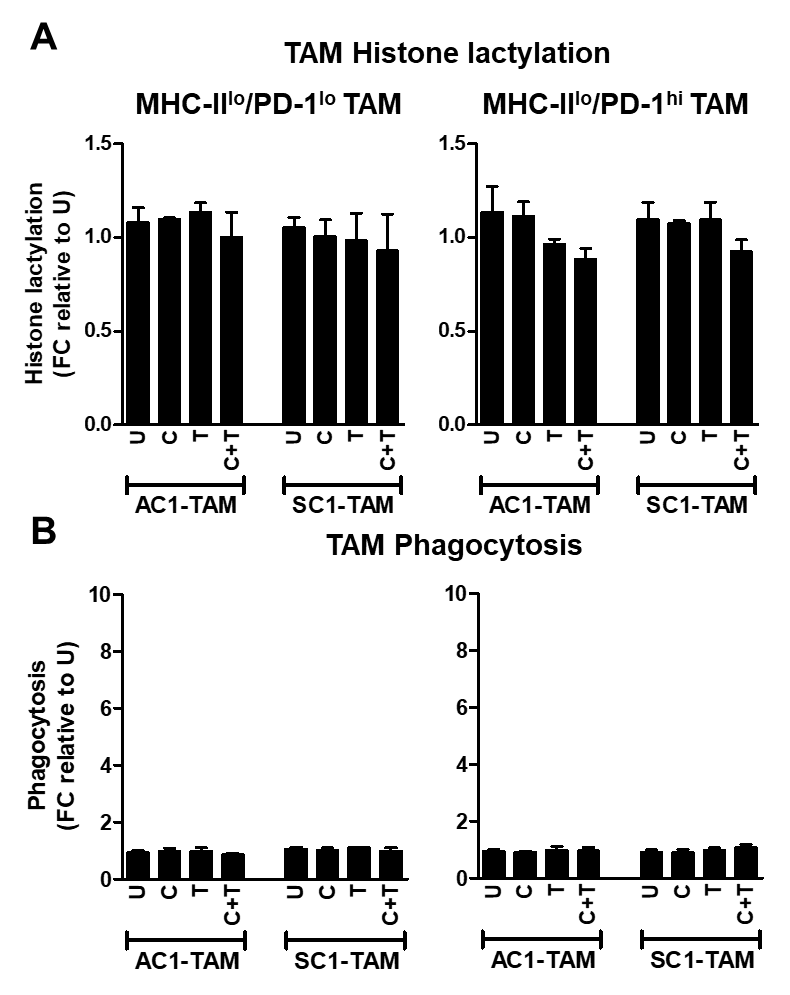

**Supplementary Figure S3. Conditioned media from PI3Ki/MEKi-treated PC cells does not alter MHC-II^lo^ TAM histone lactylation and phagocytosis.** (A-B) Single cell suspensions of PTEN/p53-deficient prostate GEM tumors were treated with copanlisib (C, 100 nM), trametinib (T, 5 nM) or their combination for 24 hours, and conditioned media (CM) was collected at the end of treatment. FACS-sorted TAM from untreated PTEN/p53-deficient GEM tumors were incubated in CM *ex vivo* for 24 hours followed by co-culture with CTV dye stained-AC1/SC1 cells for 2 hours. Bar graphs demonstrate histone lactylation status (A) and phagocytic activity (B) of MHC-II^lo^/PD-1^hi/lo^ expressing TAM, relative to untreated group. FC = fold change. N=2 independent experiments. Significances/p-values were calculated by one-way ANOVA.

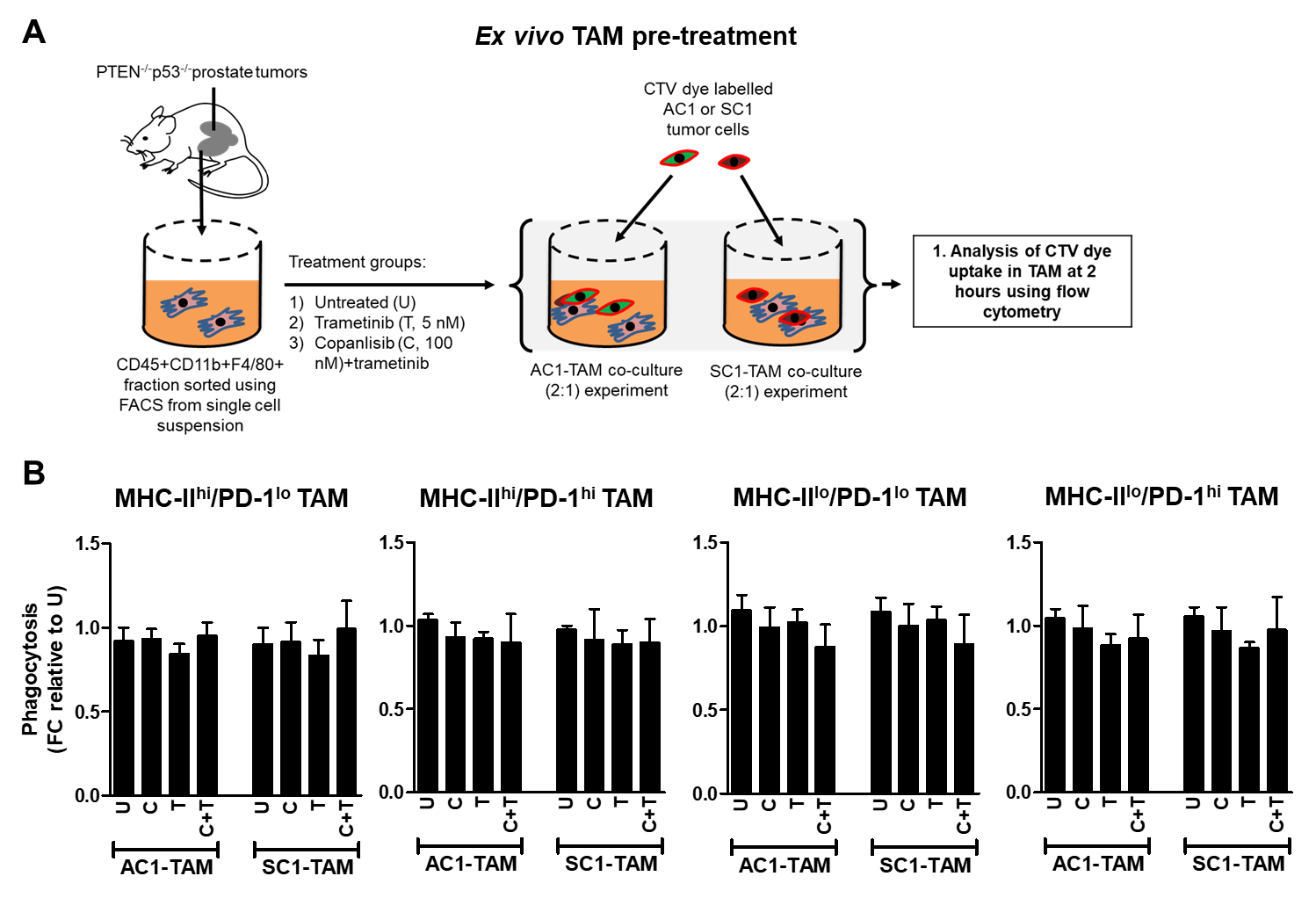

**Supplementary Figure S4. Direct treatment of TAM with PI3Ki/MEKi does not alter their phagocytic capacity.** (A) Schema illustrating direct treatment of FACS-sorted TAM subsets (PD-1^hi/lo^MHC-II^hi/lo^) from PTEN/p53-deficient prostate GEMM tumors with copanlisib (C, 100 nM), trametinib (T, 5 nM) or their combination for 24 hours. Following treatment, TAM were co-cultured with CTV dye stained-AC1/SC1 cells for 2 hours, and phagocytosis quantification was performed. (B) Bar graphs demonstrate fold change (FC) phagocytic activity of MHC-II^hi/lo^/PD-1^hi/lo^ expressing TAM, relative to untreated group. N=2 independent experiments. Significances/p-values were calculated by one-way ANOVA.

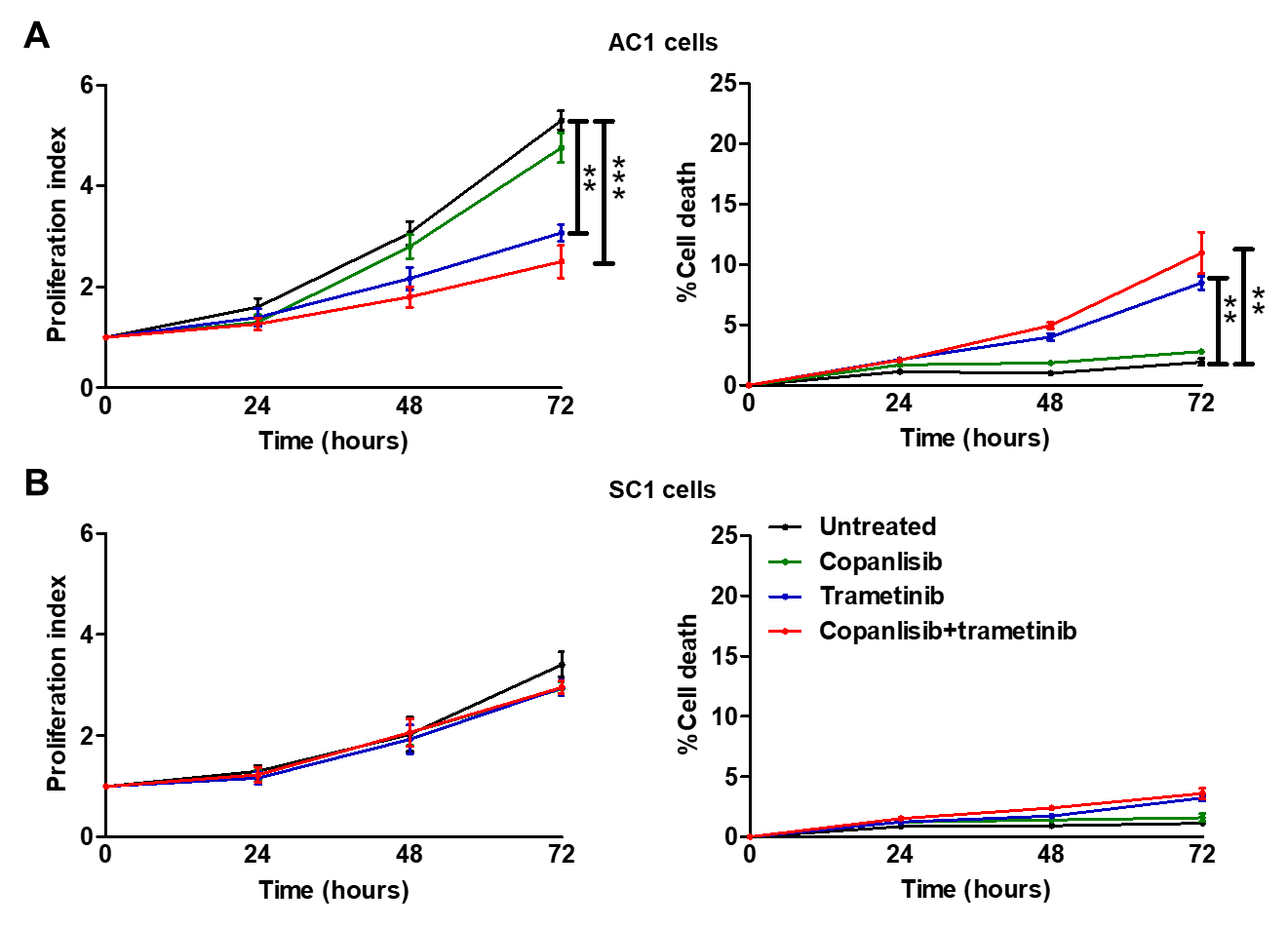

**Supplementary Figure S5. MEKi has modest anti-proliferative and pro-apoptotic effects on AC1 cells, but not SC1 cells.** (A-B) PTEN/p53-deficient GEM tumor derived cancer cell lines AC1//SC1 cells were treated with copanlisib (IC_90_ = 100 nM), trametinib (IC_90_ = 5 nM) or their combination for 24, 48 and 72 hours. (A) Proliferation index was calculated by counting number of tumor cells in each well, relative to their baseline seeding density. (B) Tumor cells were stained for annexin V antibody/propidium iodide and % cell death was analyzed by flow cytometry. n=3 independent experiments. Significances/p-values were calculated by one-way ANOVA and indicated as follows, **p<0.01 and ***p<0.001.

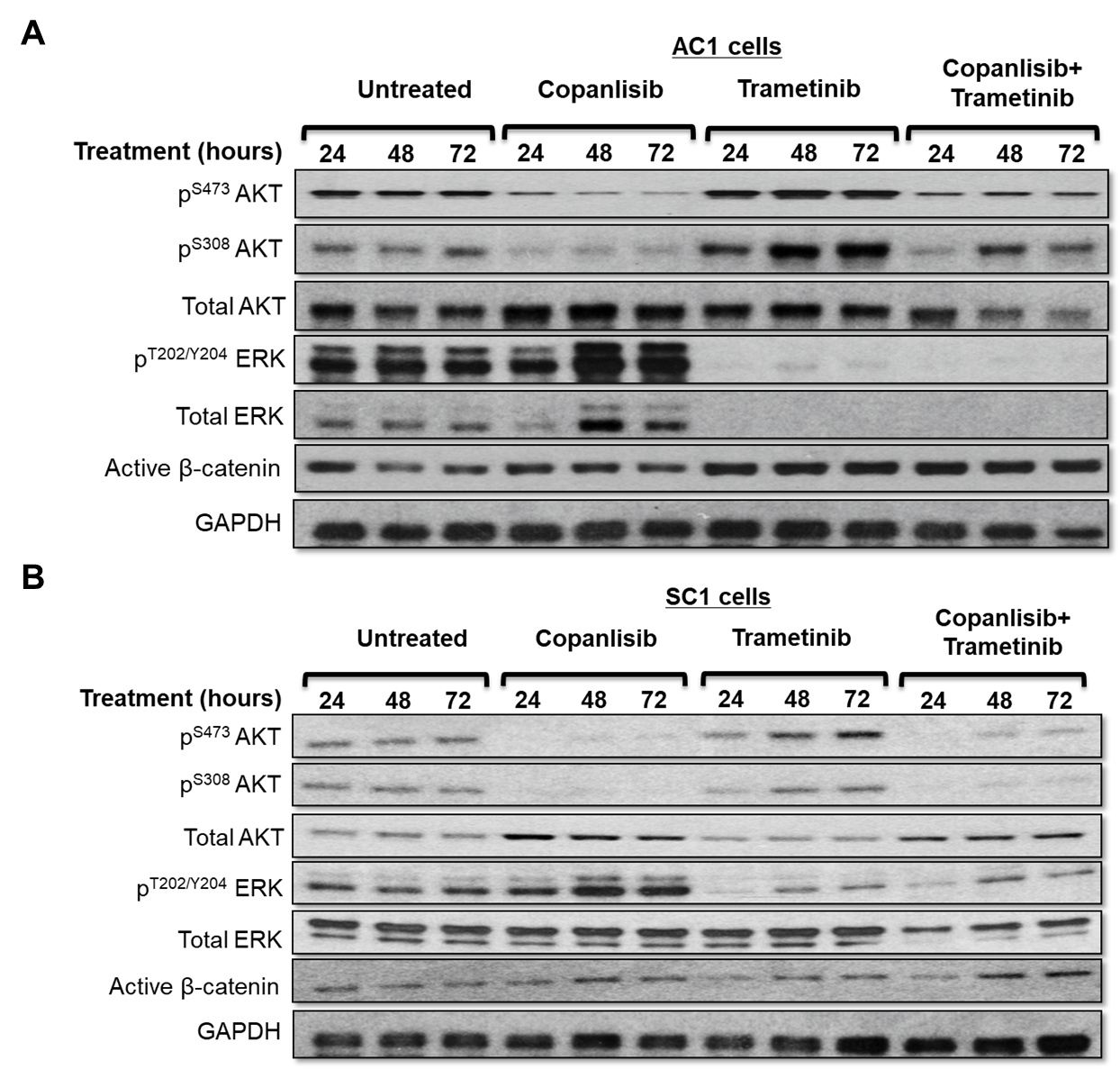

**Supplementary Figure S6. PI3Ki/MEKi combination therapy for 72 hours activates Wnt/β-catenin pathway in PTEN/p53-deficient prostate GEMM tumor derived cell lines.** (A-B) AC1/SC1 cells were treated with copanlisib (C, 100 nM), trametinib (T, 5 nM) or their combination for 24, 48 and 72 hours. Following treatment, western blot analyses with indicated protein markers were performed on AC1 (A) and SC1 (B) cell lysates. n=3 independent experiments.

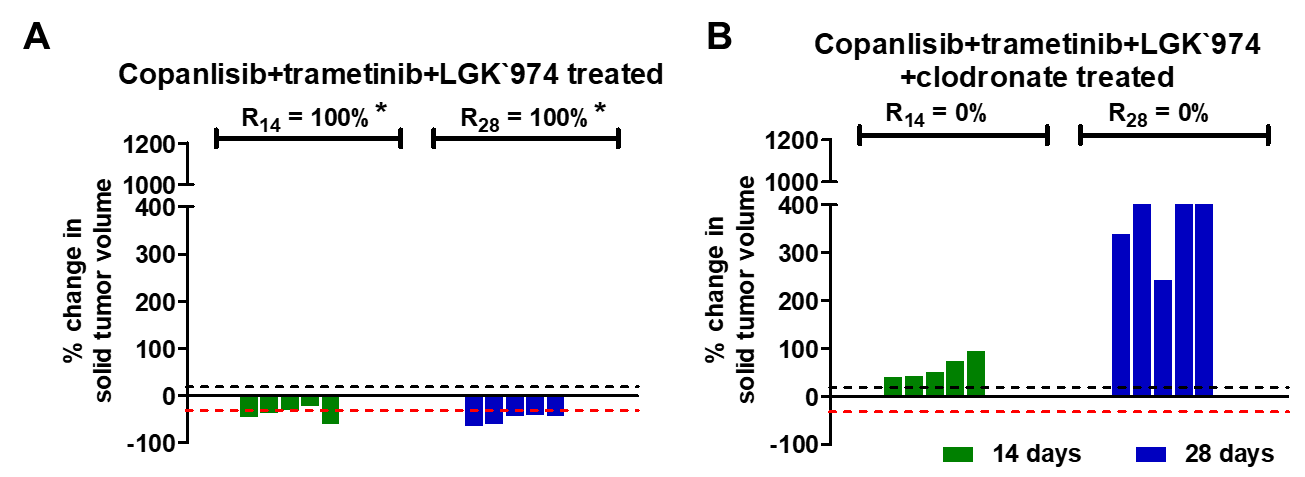

**Supplementary Figure S7. PI3Ki + MEKi + PORCNi combination therapy elicits a 100% response rate in PTEN/p53-deficient GEMMs via TAM activation.** (A-B) Pb-Cre;PTEN^fl/fl^Trp53^fl/fl^ mice with established tumors were treated with copanlisib (14 mg/kg, *iv*, every alternate day) + trametinib (3 mg/kg, *po*, every day) + LGK`974 (3 mg/kg, *po*, every day) -/+ clodronate (200 μg/mouse, *ip*, every week). Tumor volumes were non-invasively monitored by MRI and % response rate at days 14 (R_14_) and 28 (R_28_) were determined as described in Methods. The % change in solid tumor volume are represented by waterfall plot for copanlisib + trametinib + LGK`974 (A) and copanlisib + trametinib + LGK`974 + clodronate (B) treated groups. n=5 mice per group. Significances/p-values were calculated by Chi-square test and indicated as follows, *p<0.05, relative to untreated.

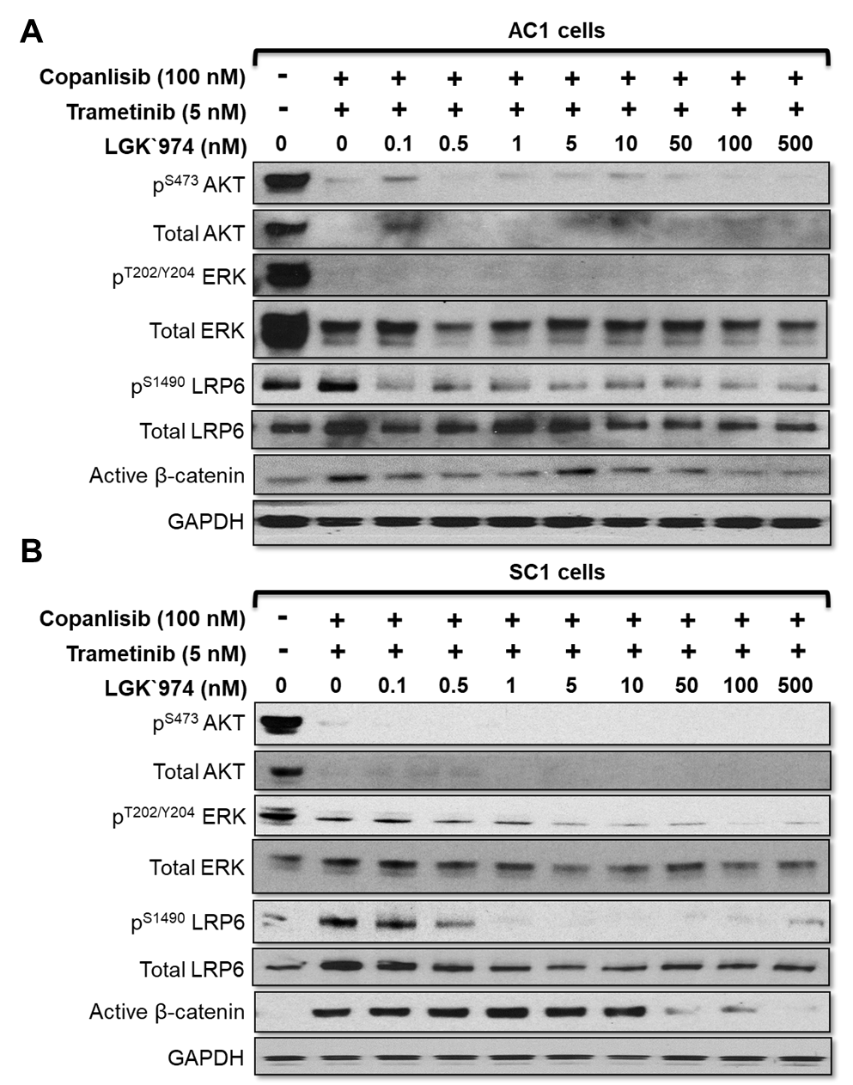

**Supplementary Figure S8. Addition of PORCNi inhibits PI3Ki + MEKi mediated Wnt/β-catenin pathway activation in PTEN/p53-deficient GEMM tumor-derived PC cells.** (A-B) AC1 (A) and SC1 (B) cells were treated with copanlisib (IC_90_ = 100 nM, 72 hours) and trametinib (IC_90_ = 5 nM, 72 hours) combination to activate Wnt/β-catenin pathway. LGK`974 (0.1-500 nM, 72 hours) was concurrently added in a dose-dependent manner and western blot analyses were performed for indicated proteins to determine IC_90_ for Wnt/β-catenin pathway inhibition. n=2 independent experiments.

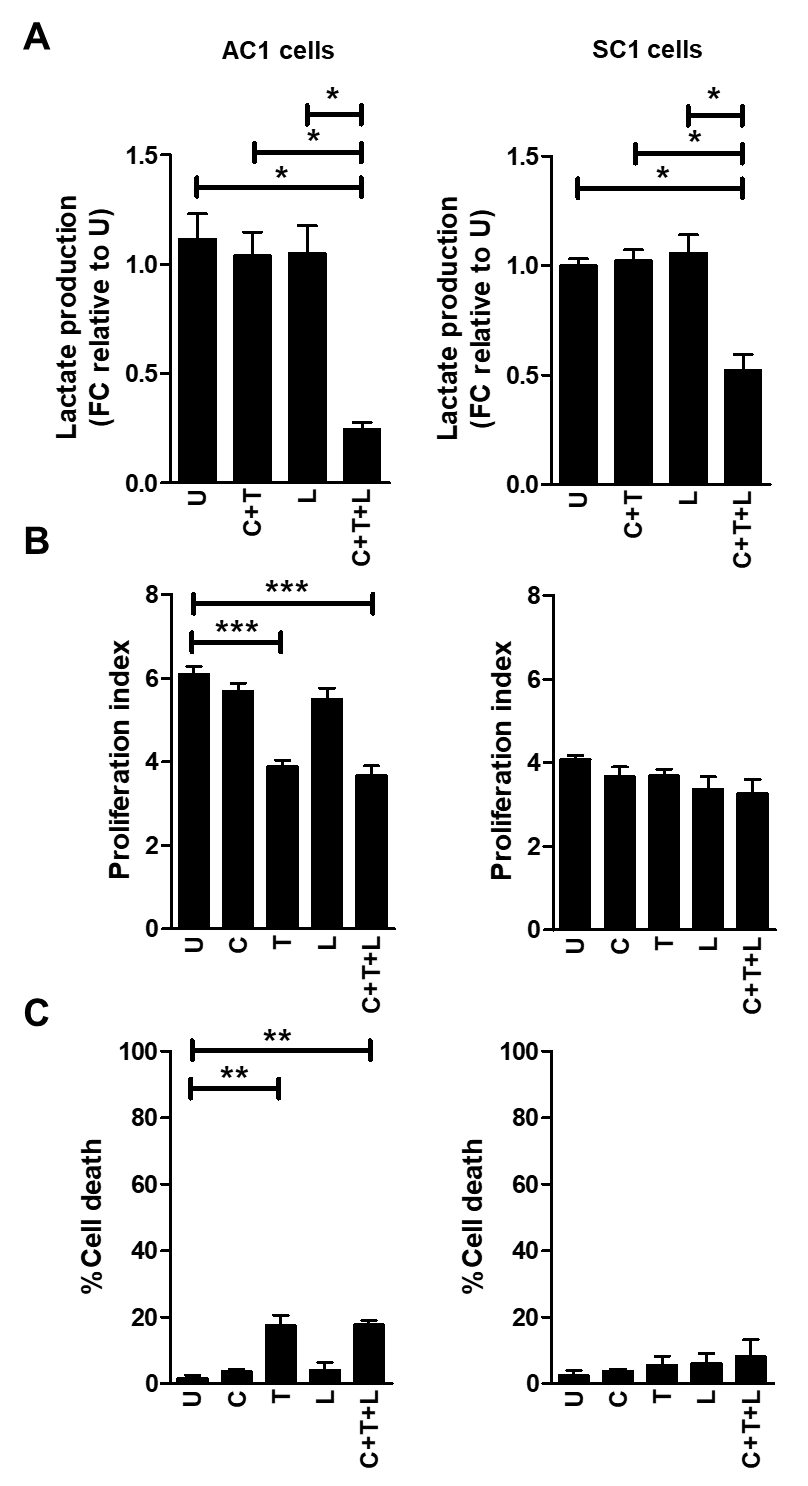

**Supplementary Figure S9. PORCNi in combination with PI3Ki + MEKi inhibits lactate production from PTEN/p53-deficient GEMM tumor-derived PC cells.** (A) AC1/SC1 cells were treated with copanlisib (C, IC_90_=100 nM), trametinib (T, IC_90_=5 nM), LGK`974 (L, IC_90_=50 nM) or their combination for 72 hours. *In vitro* CM were collected following treatment and analyzed for lactate content using colorimetry kits. (B) Proliferation index was calculated by counting number of tumor cells in each well, relative to their baseline seeding density. (C) Following treatments, tumor cells were stained for annexin V antibody/propidium iodide and % cell death were analyzed using flow cytometry. n=3 independent experiments. Significances/p-values were calculated by one-way ANOVA and indicated as follows, *p<0.05, **p<0.01 and ***p<0.001.

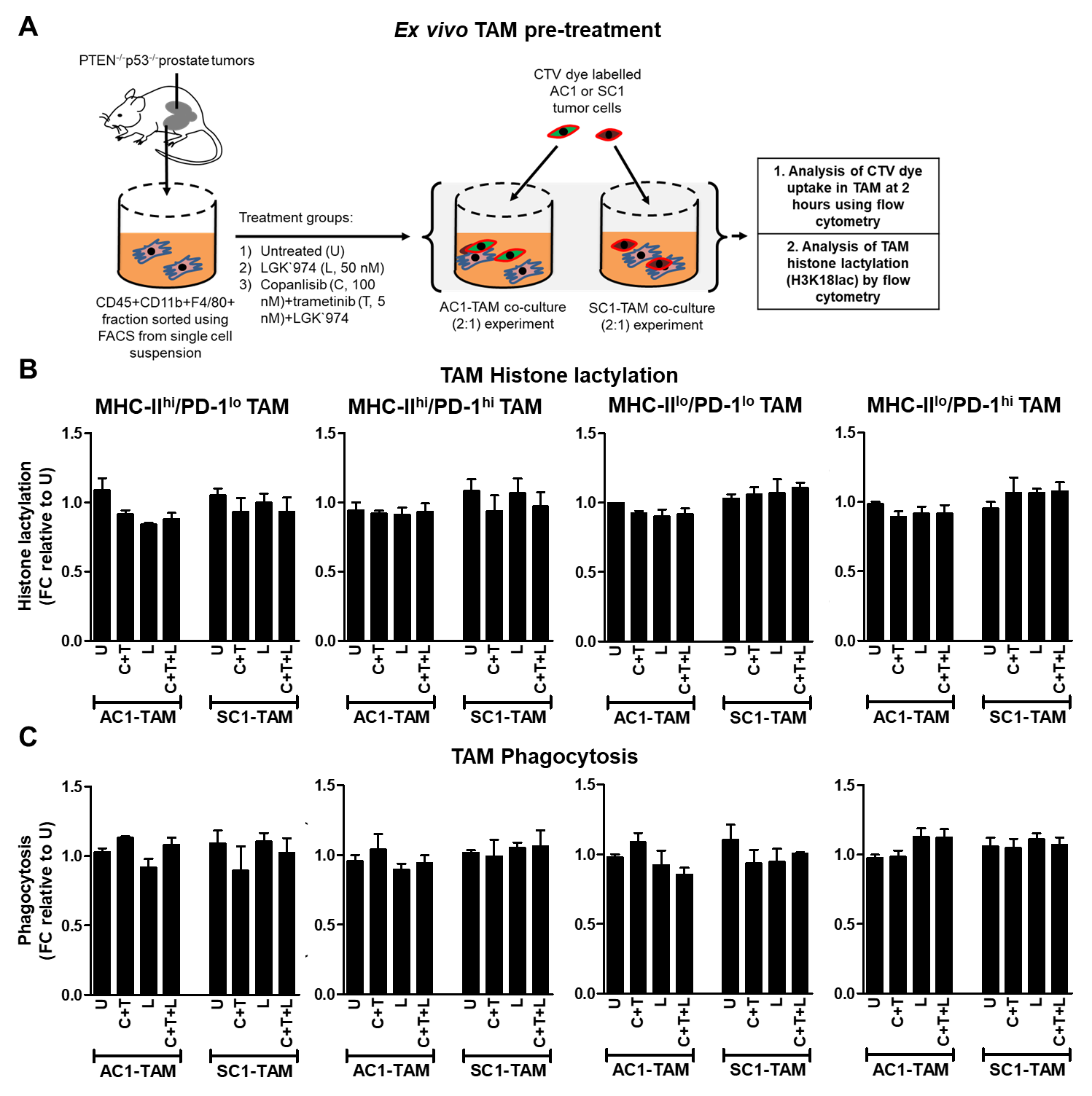

**Supplementary Figure S10. Direct treatment of TAM with PI3Ki + MEKi + PORCNi therapy does not alter TAM phagocytic capacity.** (A) Experimental schema: Direct treatment of FACS-sorted TAM subsets (MHC-II^hi/lo^/PD-1^hi/lo^) from PTEN/p53-deficient prostate tumors with copanlisib (C, 100 nM), trametinib (T, 5 nM), LGK`974 (L, 50 nM) or their combination for 24 hours. Following treatment, TAM were co-cultured with CTV dye stained-AC1/SC1 cells for 2 hours. Bar graphs demonstrate (B) histone lactylation status and (C) phagocytic activity of PD-1^hi/lo^MHC-II^hi/lo^ expressing TAM relative to untreated group. FC = fold change. n=2 independent experiments. Significances/p-values were calculated by one-way ANOVA.

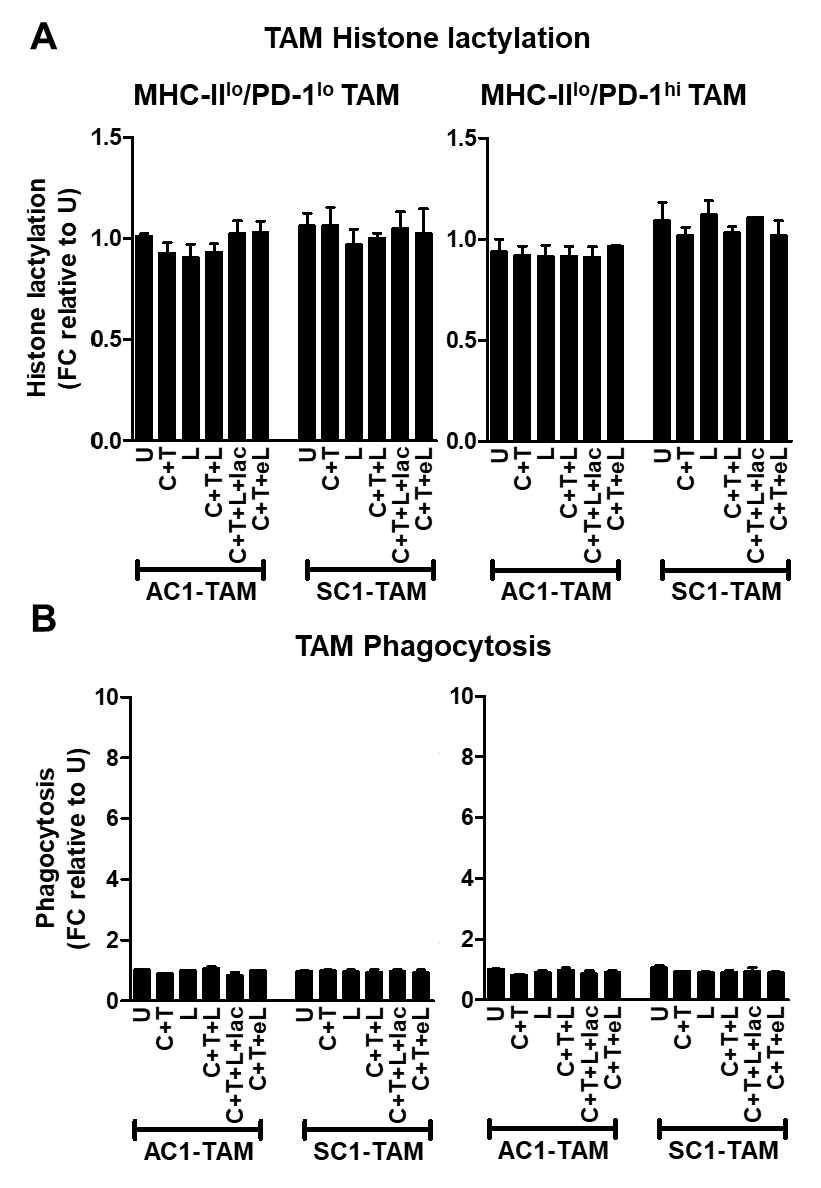

**Supplementary Figure S11. Conditioned media from PI3Ki + MEKi + PORCNi treated PTEN/p53-deficient tumor derived cells does not alter histone lactylation and phagocytic activity of MHC-II^lo^ TAM.** (A-B) AC1/SC1 cells were treated with copanlisib (C, 100 nM), trametinib (T, 5 nM), LGK`974 (L, 50 nM) or their combination for 72 hours. For mechanistic dissection, lactate (lac, 100 nmol/μL) and LGK`974 (eL, 50nM) were added to the CM collected after C+T+L and C+T treatments of AC1/SC1 cells, respectively. FACS-sorted TAM were incubated with the indicated CM for 24 hours followed by co-culture with CTV dye stained-AC1/SC1 cells for 2 hours. Bar graphs demonstrate fold change (FC) in (A) histone lactylation and (B) phagocytosis of MHC-II^lo^/PD-1^hi/lo^ expressing TAM relative to untreated group. n=2 independent experiments. Significances/p-values were calculated by one-way ANOVA and indicated as follows, ***p<0.001.

**Supplementary table S1. PI3Ki + MEKi + PORCNi therapy results in tumor clearance in Pb-Cre;PTEN^fl/fl^Trp53^fl/fl^ mice.** Pb-Cre;PTEN^fl/fl^Trp53^fl/fl^ mice with established tumors were treated with copanlisib (C, 14 mg/kg, *iv*, every alternate day) + trametinib (T, 3 mg/kg, *po*, every day) + LGK`974 (L, 3 mg/kg, *po*, every day) for 28 days and approximately 7 months, as described in Methods section. At the end of treatment, harvested tumors were formalin fixed, paraffin embedded and individual slides were stained with H&E and scored in a blinded fashion by pathologist for histopathological response, as indicated in the table.

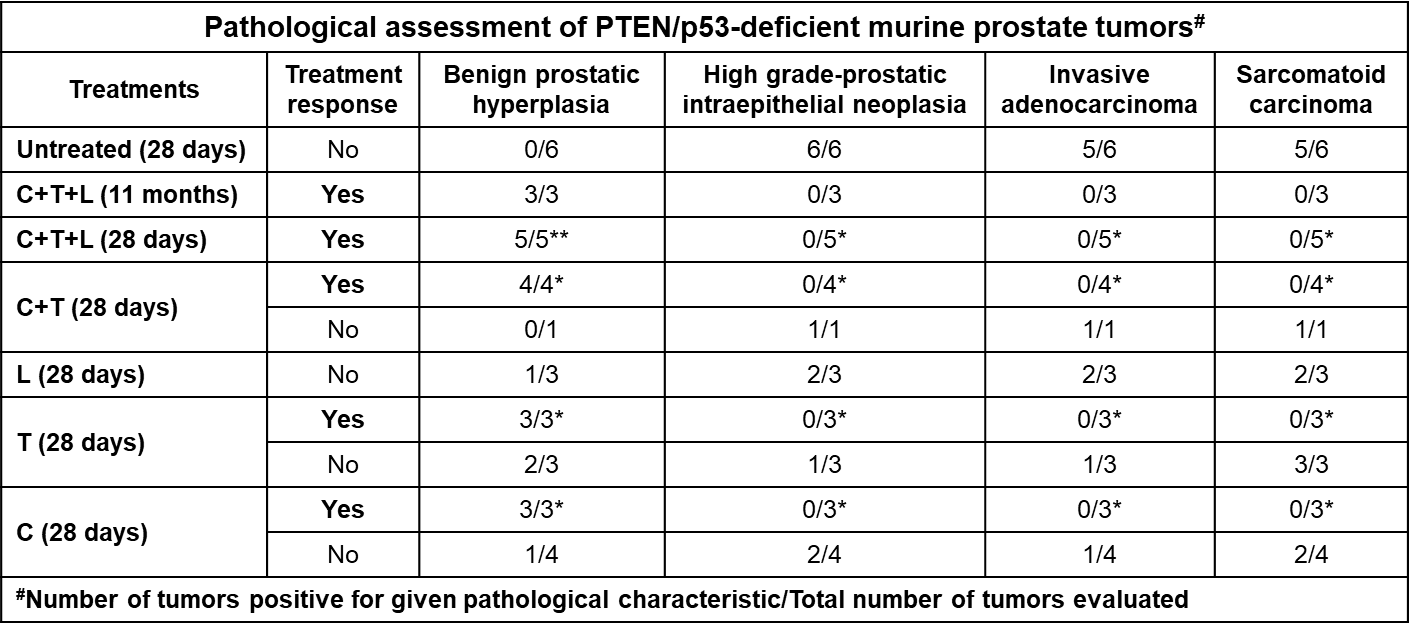
